# Evolution of a virus-like architecture and packaging mechanism in a repurposed bacterial protein

**DOI:** 10.1101/2020.12.23.423990

**Authors:** Stephan Tetter, Naohiro Terasaka, Angela Steinauer, Richard J. Bingham, Sam Clark, Andrew J. P. Scott, Nikesh Patel, Marc Leibundgut, Emma Wroblewski, Nenad Ban, Peter G. Stockley, Reidun Twarock, Donald Hilvert

## Abstract

Viruses are ubiquitous pathogens of global impact. Prompted by the hypothesis that their earliest progenitors recruited host proteins for virion formation, we have used stringent laboratory evolution to convert a bacterial enzyme lacking affinity for nucleic acids into an artificial nucleocapsid that efficiently packages and protects multiple copies of its own encoding mRNA. Revealing remarkable convergence on the molecular hallmarks of natural viruses, the accompanying changes reorganized the protein building blocks into an interlaced 240-subunit icosahedral capsid impermeable to nucleases, while emergence of a robust RNA stem-loop packaging cassette ensured high encapsidation yields and specificity. In addition to evincing a plausible evolutionary pathway for primordial viruses, these findings highlight practical strategies for developing non-viral carriers for diverse vaccine and delivery applications.

## Main Text

Understanding the origins and evolutionary trajectories of viruses is a fundamental scientific challenge.^1^ Even the simplest virions, optimized for genome propagation over billions of years of evolution, require co-assembly of many copies of a single protein with an RNA or DNA molecule to afford a closed-shell container of defined size, shape, and symmetry. Strategies for excluding competing host nucleic acids and protecting the viral genome from nucleases are also needed. While recreating such properties in non-viral containers is challenging,^2–6^ capsids generated by bottom-up design are promising as customizable tools for delivery and display.^7–9^

Previous efforts to produce artificial nucleocapsids that encapsulate their own genetic information have utilized natural and computationally designed protein cages possessing engineered cationic interiors.^5,6^ However, even after directed evolution only ~10% of the resulting particles contained the full-length target RNA, underscoring the difficulties associated with packaging and protecting nucleic acids in a cell. In addition to competition from abundant host nucleic acids, genome degradation by cellular RNases is problematic owing to slow assembly, cage dynamics and/or porosity. Here we show that complementary adaptations of cargo and container can be harnessed to address these challenges and recapitulate the structural and packaging properties of natural viruses.

Our starting point was a previously evolved nucleocapsid, derived from *Aquifex aeolicus* lumazine synthase (AaLS), a bacterial enzyme that naturally forms 60-subunit nanocontainers but has no inherent affinity for nucleic acids.^10^ AaLS was redesigned by circular permutation and appending the arginine-rich peptide λN+, which tightly binds an RNA stem-loop called BoxB^11,12^ (Supplementary Figure 1A). The resulting nucleocapsid variant, NC-1, was subsequently evolved via intermediate NC-2 to NC-3 by selecting for variants that capture capsid-encoding mRNA transcripts flanked by BoxB tags. Nevertheless, only one in eight of the NC-3 capsids packaged the full-length RNA genome.^6^

To improve NC-3’s packaging properties, we mutagenized its gene by error-prone PCR and subjected the library to three cycles of expression, purification, and nuclease challenge, followed by reamplification of the surviving mRNA. Selection stringency was steadily increased in each cycle by decreasing nuclease size (60 kDa benzonase à 14 kDa RNase A à 11 kDa RNase T1) and extending nuclease exposure from 1 to 4 hours. This strategy ensured 1) efficient assembly of RNA-containing capsids, 2) protection of the cargo from nucleases, and 3) enrichment of variants that package the fulllength mRNA (Figure 1A). The best variant, NC-4, had nine new mutations, three of which were silent (Supplementary Figure 2).

**Figure 1:**
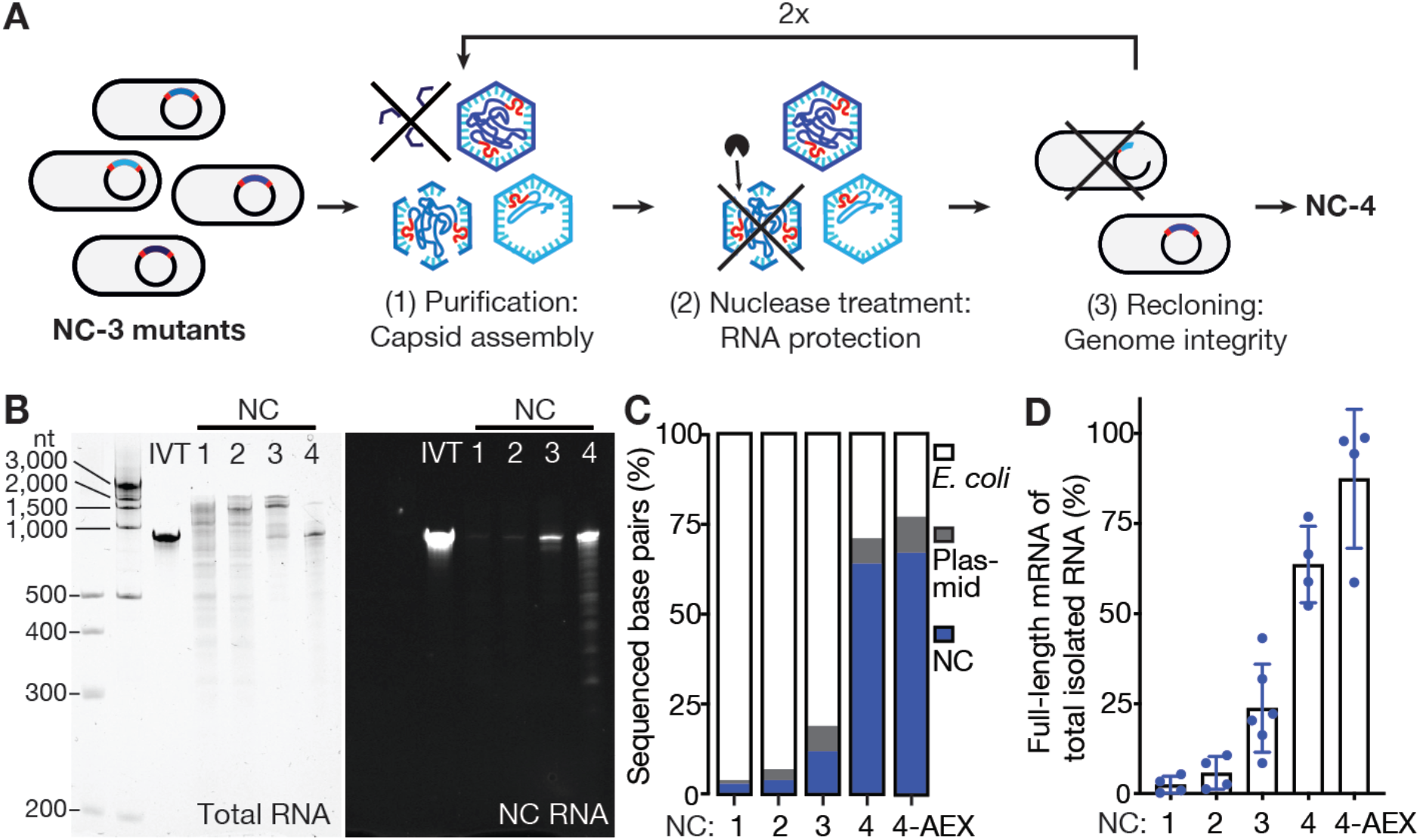
NC-4 packages its genome with high selectivity. (A) Laboratory evolution: a library of NC-3 mutants generated by error-prone PCR was expressed in Escherichia coli and purified by affinity and size exclusion chromatography. This step recovers assembled capsids. The purified capsid library was then treated with nucleases to enrich for capsids that protect their RNA cargo. Finally, the RNA was extracted from capsids, reverse-transcribed, and re-cloned into the original expression vector. This step selects for capsids that contain full-length genomes. (B) Denaturing PAGE (5%) of NC-1 to NC-4 stained for total RNA with GelRed (left) and the fluorogenic dye DFHBI-1T (right), which selectively binds the broccoli aptamer present in the 5’- and 3’-untranslated regions of the mRNA genome (NC RNA). IVT, in vitro-transcribed reference mRNA. (C) RNA identities and their relative abundance were determined by Oxford Nanopore Sequencing^33^ for all four capsids, including anion-exchanged NC-4 (4-AEX), and assigned to three main categories: bacterial RNA (E. coli), nucleocapsid mRNA (NC), and RNA originating from other plasmid-associated genes (plasmid). The encapsulated E. coli genes are primarily rRNA (Supplementary Figure 3). (D) The fraction of total extracted RNA corresponding to the full-length mRNA genome was determined by real-time quantitative PCR (mean of at least two biological replicates, each measured in two separate laboratories, error bars represent the standard deviation of the mean).

After optimizing protein production and purification, we compared NC-4 to its precursors. Particle heterogeneity decreased notably over the course of evolution from NC-1 so that NC-4 assembles into homogeneous capsids (Supplementary Figure 1B,C), with protein yields after purification (~35 mg/L medium) that increased by an order of magnitude in the last evolutionary step. Additionally, nuclease resistance steadily improved. NC-1 RNA is almost completely degraded upon treatment with either benzonase or RNase A, whereas NC-2 protects small amounts of full-length mRNA from benzonase but not RNase A (Supplementary Figure 1D). In contrast, both NC-3 and NC-4 protect most of their encapsidated RNA from both nucleases (Supplementary Figure 1D,E). Importantly, NC-4 also packages its own full-length mRNA with improved specificity. While earlier generations encapsidate a broad size range of RNA species (400–2000 nt), NC-4 binds one major species corresponding to the 863 nt-long capsid mRNA (Figure 1B, left). Long-read direct cDNA sequencing confirmed the decrease in encapsidated host RNA (Figure 1C), which was largely ribosomal (Supplementary Figure 3). The simultaneous increase in genome packaging efficiency over the four generations is clearly evident in gels stained with the fluorogenic dye DFHBI-1T, which binds the Broccoli aptamers^13^ introduced with the BoxB tags^6^ (Figure 1B, right).

The fraction of full-length genome relative to total encapsidated RNA was quantified by real-time PCR to be (2±2)% for NC-1, (6±5)% for NC-2, (24±12)% for NC-3 and (64±11)% for NC-4 (Figure 1D). When NC-4 was further purified by ion-exchange chromatography to remove incomplete or poorly-assembled capsids, (87±19)% of the RNA corresponded to the full-length genome. Given the total number of encapsidated nucleotides (~2500), NC-4 packages on average 2.5 full-length mRNAs per capsid, a dramatic improvement compared to its precursors and other artificial nucleocapsids.^5,6^ This packaging capacity suggests that the evolved capsid could readily accommodate substantially longer RNAs, such as more complex genomes or large RNA molecules of medical interest.

Improved genome packaging and protection were accompanied by major structural transformations. The cavity of the starting 16 nm diameter AaLS scaffold is too small to package an 863 nt-long RNA.^2,6^ However, addition of the λN+ peptide to circularly permuted AaLS afforded expanded capsids with diameters in the 20–30 nm range, which were subsequently evolved toward uniform ~30 nm diameter particles (Supplementary Figure 1C). To elucidate the nature of these changes, we turned to cryo-EM.

Characterization of the initial NC-1 design revealed a range of assemblies of varying size and shape (Supplementary Figure 4A,B). Although particle heterogeneity and aggregation complicated singleparticle reconstruction, two expanded structures with tetrahedral symmetry were successfully obtained (Figure 2A). Like the wild-type protein, both are composed entirely of canonical lumazine synthase pentamers (Figure 3A), but they possess large, keyhole-shaped pores (~4 nm wide) through which nucleases could diffuse. One capsid is a 180-mer (Supplementary Figure 4C–F, Table S1) that closely resembles a previously characterized AaLS variant possessing a negatively charged lumen.^14,15^ The other NC-1 structure is an unprecedented 120-mer (Supplementary Figure 4G–I, Table S1). It features wild-type-like pentamer-pentamer interactions as well as inter-pentamer contacts characteristic of its 180-mer sibling (Figure 2A). At the monomer level, the major deviation from the AaLS fold is seen in a short helix (residues 67–74) and adjacent loop (residues 75–81) (Figure 2B,C). In AaLS, this region is involved in lumenal interactions between the pentameric building blocks at the threefold-symmetry axes. In NC-1 chains that are not involved in wild-type-like pentamer-pentamer contacts, this loop assumes altered conformations and is resolved to lower local resolution (Figure 2B,C, Supplementary Figure 4E,H).

**Figure 2:**
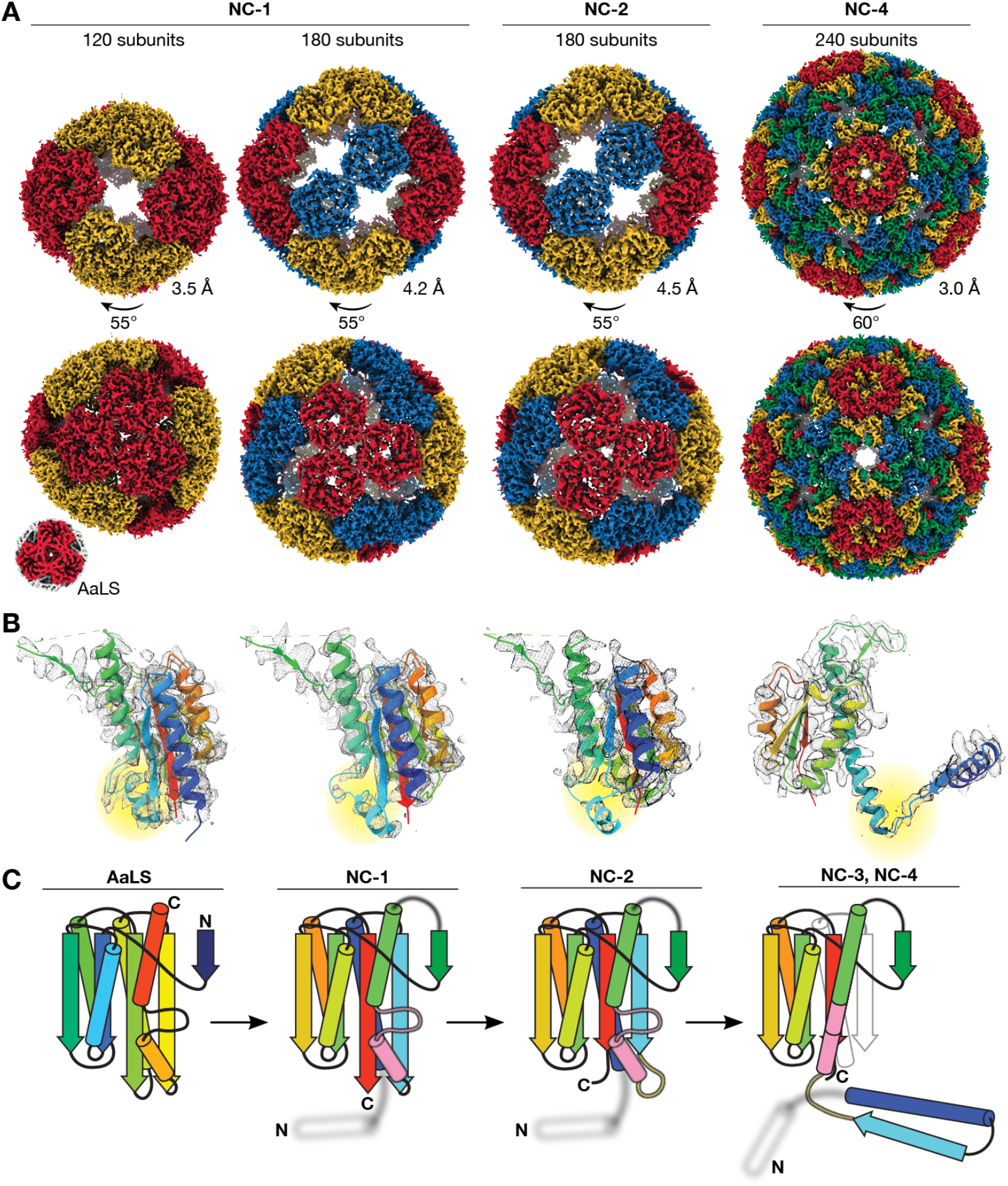
Structural evolution towards virus-like nucleocapsids. (A) Maps are shown for the tetrahedrally symmetric NC-1 and NC-2 structures with symmetry-related pentamers in the same color, and the icosahedral *T=4* NC-4 capsid with the four quasi-equivalent chains highlighted in different colors. The lower resolution NC-3 capsid (7.0 Å, Supplementary Figure 6) resembles NC-4. Wild-type AaLS^10^ is shown for comparison (not to scale). Resolutions were estimated by Fourier shell correlation (0.143 threshold). (B) Fits of single chains (rainbow; N-terminus to C-terminus from blue to red) in the electron density of the capsids above show the evolution of the monomer. Residues 66 to 81 are highlighted (yellow). Clear density is seen for this segment in NC-1 protomers involved in AaLS-like inter-pentamer contacts. In other chains, as in the 180-mer NC-1 structure, this region is less well resolved. In NC-2 the nearby beta-sheet is also perturbed, further enhancing the flexibility of this region. In NC-3 and NC-4, this segment rearranges into an extended helix that supports the domain swap. (C) Rainbow-colored models depict the changes in the protein fold. The helix (67–74) and loop (75–81) that undergo a major rearrangement are colored in pink, and the hinge loop (62–66) in yellow; the structurally unresolved RNA-binding peptide is depicted as a blurry white helix.

**Figure 3:**
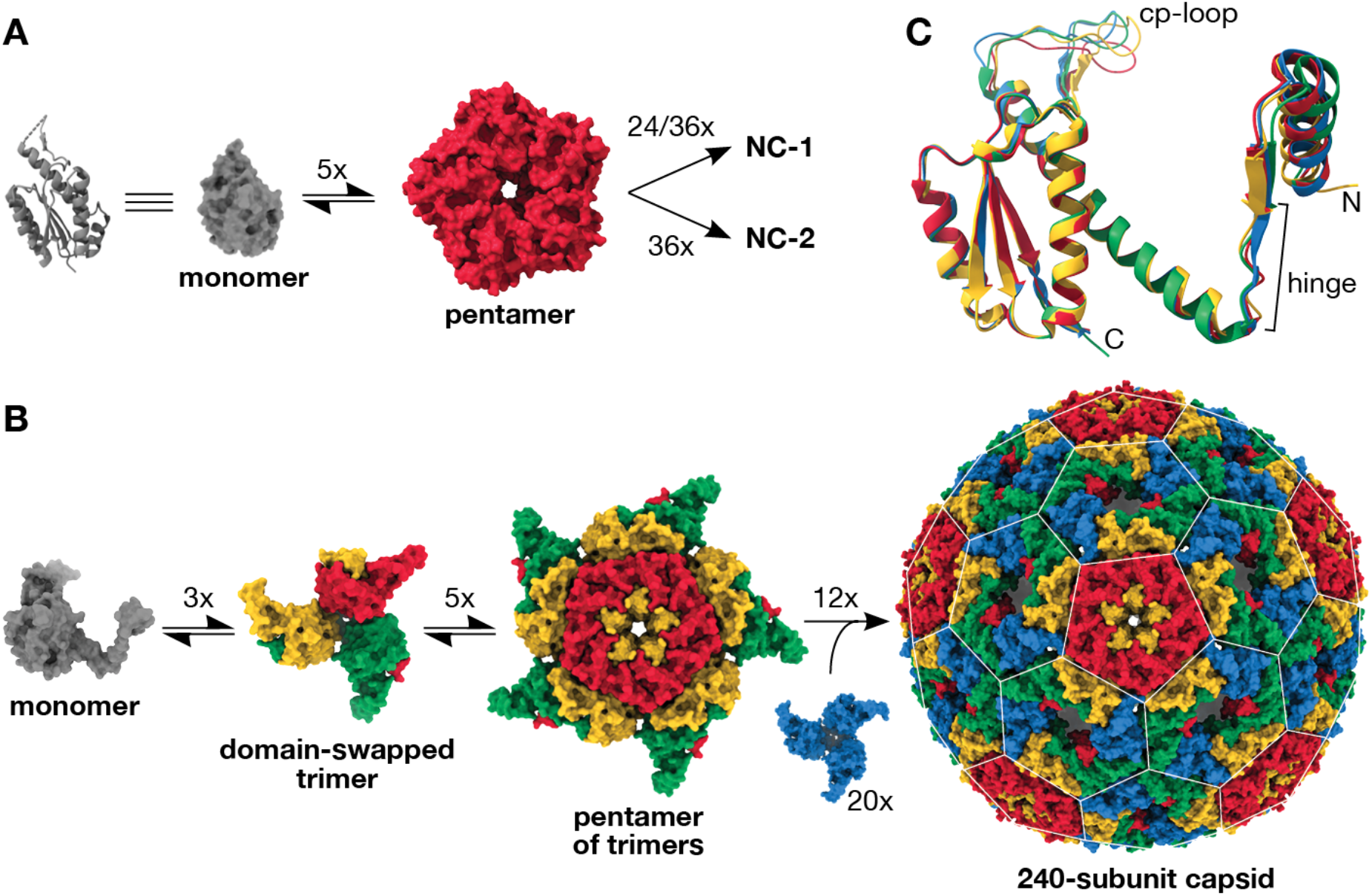
Virus-like architecture by protomer reorganization. (A) The assembly of 120- and 180-subunit NC-1 and NC-2 cages from monomers (cartoon and surface shown in grey) presumably proceeds via AaLS-like pentamers. (B) Based on the assembly mechanisms of other viral capsids,^34^ the T=4 capsids likely arise from domain-swapped trimeric building blocks that further combine into pentamers. Combining the latter with additional domain-swapped trimers (blue) would afford the complete 240-subunit capsid. The pentagonal and hexagonal faces of the icosahedrally symmetric capsid are highlighted by a white lattice. (C) Assembly of the T=4 icosahedral NC-3 and NC-4 structures requires the subunits to adopt different, quasi-equivalent conformations. An overlay of the four quasi-equivalent chains of NC-4, colored as in panel b, shows that the hinge region provides flexibility for subtle adjustments in the relative orientation of the flanking segments. Additional differences are visible in the poorly resolved surface loop introduced by circular permutation (cp-loop), which interacts with the neighboring subunit in both pentamers and hexamers via a single short beta strand.

The second-generation variant NC-2, obtained after benzonase challenge, is also polymorphic and aggregation-prone. Several distinct morphologies were identified by 2D-classification (Figure S5), one of which was reconstructed as a tetrahedrally symmetric 180-mer (4.5 Å, Figure 2A, Table S1) that superimposes on the analogous NC-1 structure. Four mutations (I58V, G61D, V62I, and I191F) shorten two strands of the core beta-sheet and, indirectly, further increase disorder in neighboring residues 66– 81 (Figure 2B,C). These changes likely disfavor wild type-like pentamer-pentamer interactions, explaining the absence of smaller capsids with more tightly packed capsomers. Structural heterogeneity and particle aggregation precluded reconstruction of additional structures that may contribute to the benzonase-resistance phenotype.

The ability of NC-3 and NC-4 to protect their cargo from RNases significantly smaller than the pores in the parental structures suggests a novel solution to nuclease resistance. In fact, three-dimensional reconstructions of NC-3 (7.0 Å) and NC-4 (3.0 Å) (Supplementary Figure S6, Table S1) yielded superimposable structures that are markedly different from any previously characterized AaLS derivative (Figure 2A). Both capsids form icosahedrally symmetric 240-mers that feature smaller pores (~2.5 nm) than their progenitors. The pentagonal vertices align with AaLS pentamers, and are surrounded by 30 hexagonal patches (Figure 3B). This architecture is typical of T=4 virus capsids, in which a single protein chain assumes four similar, quasi-equivalent conformations, repeated with icosahedral symmetry to afford a closed container with increased volume.^16^

The most striking feature of our evolved cages is a 3D-domain swap,^17^ which links neighboring monomers and reorganizes the structure into trimeric building blocks (Figure 3B). As reported for some viral capsids,^18–21^ such interlacing may enhance particle stability. This rearrangement was made possible by a hinge around residues 62–66, which permits dissociation of the N-terminal helix and strand of each subunit from the core, allowing it to dock onto a neighboring subunit in the trimeric capsomer. An elongated alpha-helix extends C-terminally from this hinge, formed by fusing the short helix (residues 67–74) to the following helix by ordering of the intervening loop (residues 75–81) (Figure 2C). Slight variations in the hinge angles allow the subunits to occupy four quasi-equivalent positions within the expanded icosahedral lattice^22^ (Figure 3C). Such flexible elements might similarly be exploited for the rational design of large (T>1) capsid assemblies from a single protein chain, an as yet unmet challenge due to the difficulty of designing proteins capable of adopting several distinct conformations.

The smaller pores in the NC-3 and NC-4 shells provide a compelling explanation for nuclease resistance. The structurally unresolved lN+ peptides, which line the lumenal edge of these openings, likely further restrict access to the cage interior. Nevertheless, the superimposable structures do not account for the differences in packaging efficiency between NC-3 and NC-4. Although a lysine to arginine mutation that appeared in the lN+ peptide of NC-4 is known to increase affinity to the BoxB tags ~3-fold,^11^ the effects of reverting this mutation are modest (Supplementary Figure 7), indicating that other factors must be at play.

Some viruses that package single-stranded RNA genomes utilize multiple stem-loop packaging signals to ensure cargo specificity and orchestrate capsid assembly within the crowded confines of the cell.^23^ Could the evolution of additional RNA packaging signals in the NC-4 genome explain its superiority to NC-3? Besides the originally introduced BoxB tags,^6^ BB1 and BB2, both genomes have 37 BoxB-like URxRxRR (R = purine) and URxR sequences^24^ (Table S2). In order to determine whether any of these serve as packaging signals, we used synchrotron X-ray footprinting (XRF). Synchrotron radiation generates hydroxyl radicals, which cleave the RNA backbone. Because base-pairing and contact with protein decrease local cleavage propensity, XRF provides a means to map intermolecular interactions and RNA secondary structure.^25^

Footprints for packaged NC-3 and NC-4 RNA show that only BB1, BB2, and 11 out of 37 BoxB-like motifs exhibit low XRF reactivity (Table S2). Furthermore, XRF-informed prediction of RNA secondary structure ensembles^26,27^ indicates that only seven of these motifs (BB1, BB2, and potential packaging signals PS1–5) are presented as stem-loops with significant frequency (Supplementary Figures 8A, 9A). Assuming that interactions with the λN+ peptides stabilize the stem-loops, comparison of their display frequency in encapsulated versus free RNA pinpoints which of these motifs might serve as packaging signals.

In NC-3, the secondary structure predictions (Figure 4A,C,E, Supplementary Figure 8) indicate that the original high-affinity BoxB tags are more frequently displayed as stem-loops in free transcripts than in capsids (96% vs. 63% for BB1 and 75% vs. 52% for BB2). Although the five lower affinity PS1–5 motifs are displayed more frequently upon encapsulation, their broad distribution, coupled with modest display of the high-affinity tags, contrasts with natural viruses, which appear to utilize narrow clusters of packaging signals surrounding an efficiently displayed, high-affinity stem-loop to initiate capsid assembly.^23^ The lack of robust assembly instructions may explain why 72% of the RNA packaged in NC-3 is ribosomal. Ribosomal RNA is compact, abundant and also possesses multiple BoxB-like signals (Supplementary Figure 3C,D), which may allow it to function as an alternative nucleation hub for capsid assembly.

**Figure 4:**
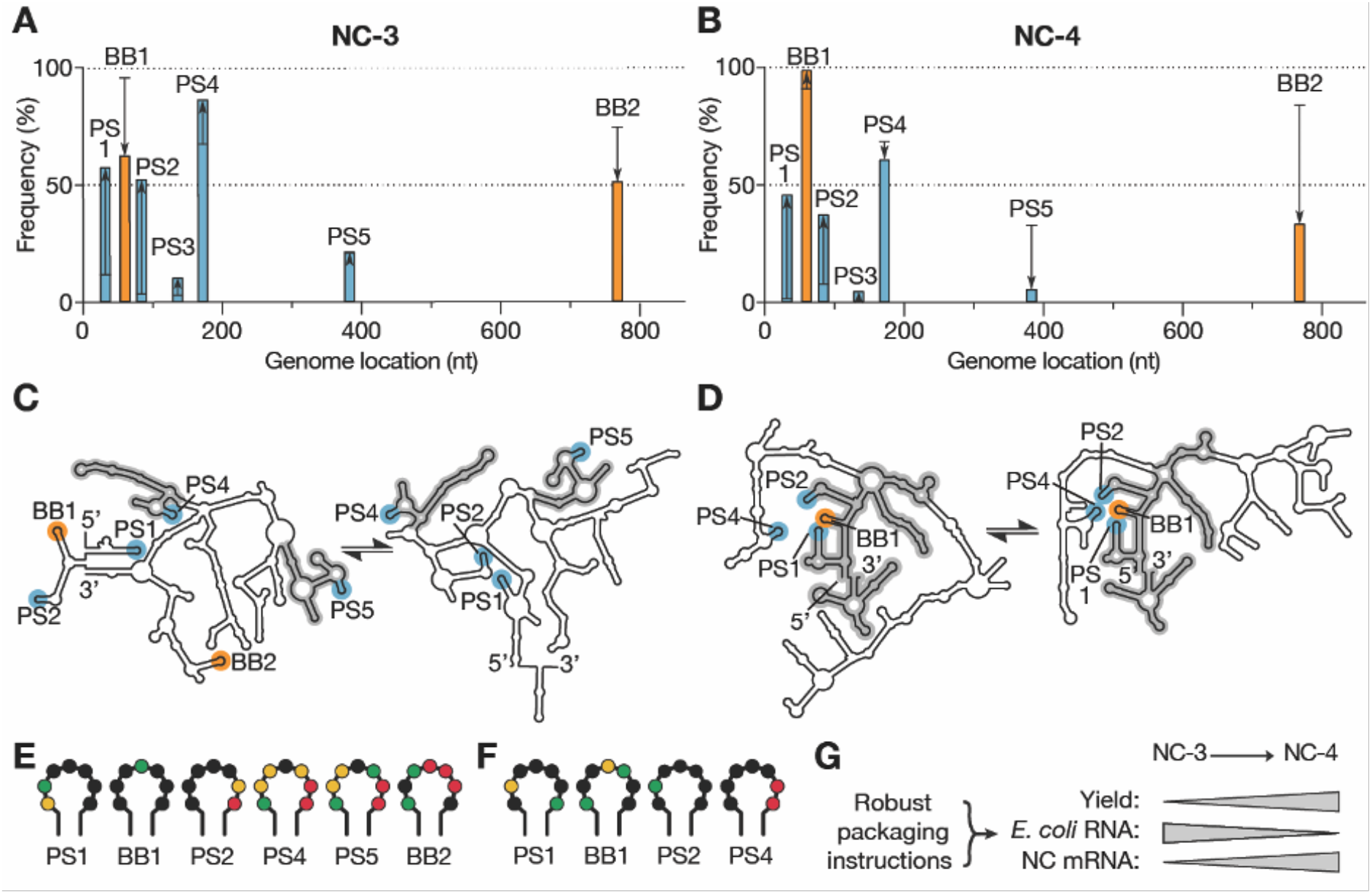
Virus-like genome packaging mediated by packaging signals. (A,B) XRF reactivities were used to calculate how frequently the seven packaging signal candidates occur in stem-loops in the NC-3 (A) and NC-4 (B) mRNA genomes; 1000 sample folds were generated for each of 1116 combinations of reactivity offsetting and scaling factors (see Supplementary Figures 8,9). Cumulative display frequencies of the motifs as stem-loop are plotted against genome position for the packaged transcripts (bars), with the high-affinity BoxB tags highlighted in orange, BoxB-like PS1–5 motifs in blue, and the respective in vitro-transcribed RNA indicated by black lines; arrows show the increase or decrease observed upon packaging. (C,D) Two consensus folds predicted for packaged NC-3 and NC-4 mRNA (see also Supplementary Figures 8,9). Secondary structure features shared between the respective folds are highlighted in grey. These structures indicate more extensive fold conservation in NC-4, as well as more robust display of a packaging cassette comprising PS1, BB1, PS2, and PS4, than in NC-3. (E,F) Reactivities of the URxRxRR motifs displayed in the packaged NC-3 (E) and NC-4 (F) RNA folds depicted in panels (C) and (D), respectively. Reactivity follows the order: red (high)à yellowà greenà black (low). The four packaging signal candidates in NC-4 show low reactivities, consistent with protection by capsid protein. (G) The evolution of a packaging cassette that steers efficient capsid assembly around the target RNA provides a compelling explanation for the improved properties of NC-4 compared to NC-3.

In NC-4, four of the seven potential packaging signals are significantly populated as stem loops in packaged genomes (PS1, BB1, PS2, PS4) and all are clustered at the 5’-end of the transcript. Notably, BB1 is displayed in 99% of all packaged RNA folds (Figure 4B,D, Supplementary Figure 9). The low reactivities observed for the four URxR sub-motifs within the capsid (Figure 4F) imply that they are in contact with protein. Robust display of a high-affinity packaging signal within a cassette of lower affinity motifs (PS1, PS2, and PS4) is reminiscent of nucleation complexes found in Satellite Tobacco Necrosis Virus,^28^ MS2 phage,^29^ and Hepatitis B Virus.^30^ This finding suggests that NC-4 similarly evolved a key hallmark of RNA packaging signal-mediated assembly. Genome-encoded packaging instructions likely foster selective RNA encapsulation as well as rapid, efficient capsid assembly,^31^ providing a compelling explanation for the improved properties of the evolved cage (Figure 4G). Encapsulation of alternative or longer, more complex genomes may similarly benefit from optimization of RNA sequence and structure.

Successful conversion of a bacterial enzyme into a nucleocapsid that packages and protects its own encoding mRNA with high efficiency and selectivity shows how primordial self-replicators could have recruited host proteins for virion formation.^1^ The convergence on structural properties characteristic of natural RNA viruses through co-evolution of capsid and cargo is striking. Introduction of destabilizing mutations into the starting protein was key to the dramatic remodeling of the protein shell, providing the molecular heterogeneity needed to depart from the initial, energetically stable, architectural solution and converge on a regular, 240-subunit, closed-shell icosahedral assembly. At the same time, evolution of multiple RNA packaging motifs that can cooperatively bind the coat proteins likely guided specificity and efficient assembly. While such constructs are themselves attractive as customizable and potentially safe alternatives to natural viruses for gene delivery and vaccine applications, the lessons learned from their evolution may also inform ongoing efforts to tailor the properties of natural viruses for more effective gene therapy. ^32^

## Acknowledgements

We acknowledge technical support from the Functional Genomics Center Zurich, Miroslav Peterek and Peter Tittmann (Scientific Center for Optical and Electron Microscopy, ETH Zurich), Daniel Böhringer (Cryo-EM Knowledge Hub, ETH Zurich) for help with cryoEM Data collection and elaboration, and Oliver Allemann for help with optimization of nucleocapsid expression and purification. We thank DNA Sequencing & Services (MRC I PPU, School of Life Sciences, University of Dundee, Scotland, www.dnaseq.co.uk) for DNA sequencing. DH thanks the Swiss National Science Foundation and the ETH Zurich for financial support. RT acknowledges funding via an EPSRC Established Career Fellowship (EP/R023204/1) and a Royal Society Wolfson Fellowship (RSWF/R1/180009). RT & PGS acknowledge support from a Joint Wellcome Trust Investigator Award (110145 & 110146), and PGS also thanks the NSLS-II at the Brookhaven Synchrotron for the award of slots for data collection at Beamline 17-BM, together with Erik Farquhar for his expert assistance during these visits. NT acknowledges support from the Human Frontier Science Program (LT000426/2015-L). AS is the recipient of a Marie Skłodowska-Curie Individual Fellowship (LEVERAGE mRNA).

## Author contributions

NT performed laboratory evolution, AS, ST, and NT biochemical characterization of nucleocapsids, and ST cryo-EM analysis with help from ML. APS, NP, and EW performed, and RJB, SC, PGS, and RT analyzed X-ray footprinting experiments. ST, AS, and DH wrote the manuscript with input from all other authors.

## Data availability

The principal data supporting the findings of this study are provided in the figures and Supplementary Information. Additional data that support the findings of this study are available from the corresponding author upon request. Cryo-EM maps and atomic models have been deposited in the Electron Microscopy Data Bank (EMDB) and wwPDB, respectively, with the following accession codes: EMDB-11631 and PDB 7A4F (NC-1 120-mer), EMDB-11632 and PDB 7A4G (NC-1 180-mer), EMDB-11633 and PDB 7A4H (NC-2), EMDB-11634 and PDB 7A4I (NC-3), and EMDB-11635 and PDB 7A4J (NC-4).

