## Supplementary Information for "Evolution of a virus-like architecture and packaging mechanism in a repurposed bacterial protein"

#### Methods

The nomenclature for the previously published nucleocapsids[1] was simplified to make the evolutionary relationship between the different variants clearer. NC-1 corresponds to λcpAaLS, the original nucleocapsid generated by circular permutation of AaLS and addition of the λN+ peptide to the new lumenal N-terminus; NC-2 is the best variant obtained after one round of optimization, λcpAaLS-β16; and NC-3 is the best performing variant from the second evolutionary round, λcpAaLS-α9.[1] The final variant, NC-4, was evolved in the current study.

Library construction by error-prone PCR

Error-prone PCR was carried out using the JBS Error-Prone Kit (#PP-102, Jena Bioscience) according to the manufacturer's protocol. Two primers (primer 1: 5'- GCG GAT AAC AAT TCC CCT CTA GAG; primer 2: 5'- GGG TTA TGC TAG TTA TTG CTC AGC G) were used with pMG-dB-λcpAaLS-α9 (ref. 1) as a template. PCR products were purified using the Zymoclean Gel DNA Recovery Kit (#D4001, Zymo Research). Both the products and the acceptor vector (pMG-dB) were doubly digested at their NdeI and XhoI restriction sites. The DNA fragments were purified using the DNA Clean & Concentrator-5 (#D4013, Zymo Research), ligated with T4 DNA ligase (#M0202, NEB), and purified again using the same kit. The capsid library (~1 µg ligation product) was transformed into electrocompetent Escherichia coli XL1-Blue cells by electroporation. The cells were incubated in 50 mL Luria-Bertani (LB) medium for 1 hour at 37 °C. The library size (~3 x $10^6$ mutants) was determined by plating serial dilutions of the cell suspension onto LB-agar plates containing ampicillin (50 µg/mL). To the remaining cells, LB medium and ampicillin (50 µg/mL) were added to the original volume of 50 mL. Cells were cultured overnight at 37 °C and 230 rpm. The next day, plasmid DNA was extracted using the ZR Plasmid Miniprep Classic Kit (#D4016, Zymo Research).

Directed evolution of NC-4 from NC-3

Evolution of NC-4 was based on a plasmid library generated by error-prone PCR as described above. The library was subjected to three iterative cycles of selection. Each cycle involved transformation of E. coli cells, expression of the nucleocapsid variants, isolation, and purification by affinity and size, nuclease treatment, RNA extraction, reverse transcription, and re-ligation of the surviving variants into the vector backbone. Nuclease selection stringency was increased in each cycle.

Initially, electrocompetent E. coli BL21(DE3)-gold cells were transformed with the plasmid library and incubated in 4 mL LB medium for 1 hour at 37 °C. After adding ampicillin (50 µg/mL), cells were cultured at 37 °C and 230 rpm for an additional 6 hours. This 4-mL culture was then transferred to 400 mL LB medium containing ampicillin (50 µg/mL) and cultured as before until the OD600 reached 0.4–0.6, at which point protein production was induced by adding isopropyl ß-D-1-thiogalactopyranoside (IPTG) to a final concentration of 0.2 mM. Cells were cultured at 20 °C and 230 rpm for 16 hours, then harvested by centrifugation at 5,000 *g* and 4 °C for 10 min. The pellet was stored at -20 °C until purification. For purification, cells were resuspended in 15 mL lysis buffer (50 mM sodium phosphate buffer, pH 7.4) containing 1 M NaCl and 20 mM imidazole supplemented with lysozyme (1 mg/mL; #A3711, AppliChem) and DNase I (10 µg/mL; #A3778, AppliChem). The mixture was incubated at room temperature for 1 hour. After lysis by sonication and clearance by centrifugation at 9,500 *g* and 25 °C for 25 min, the supernatant was loaded onto 2 mL of Ni(II)-NTA agarose resin (QIAGEN) in a gravity flow column. Beads were washed with lysis buffer containing 1 M NaCl and 20 mM imidazole, and protein was eluted with lysis buffer containing 200 mM NaCl and 500 mM imidazole. The buffer was exchanged to 50 mM sodium phosphate buffer (pH 7.4), 5 mM EDTA (storage buffer) containing 200 mM NaCl, using an Amicon Ultra-15 centrifugal filter unit (30 kDa MWCO, Merck Millipore). Capsids were further purified by size-exclusion chromatography (SEC) on a Superose 6 increase 10/300 GL (GE Healthcare) in storage buffer with 200 mM NaCl. Proteins were purified at room temperature.

For the selection of capsids that protect their RNA genome from nucleases, a solution of capsids containing approximately 2 µg of total RNA was treated at 37 °C for 1 hour with benzonase (2.5 U/µL; #101654, Merck Millipore) in 250 µL of storage buffer supplemented with 5 mM $MgCl_2$. RNA was then extracted with TRIzol reagent (#15596026, Invitrogen) and dissolved in water. The resulting RNA sample was incubated with RQ1 DNase (#M6101, Promega) in the manufacturer's reaction buffer at 37 °C for 1 hour and subsequently purified by phenol-chloroform extraction and ethanol precipitation. From this RNA, complementary DNA (cDNA) was prepared by reverse transcription with primer 3 (primer 3: 5'- GCG GAT AAC AAT TCC CCT CTA GAG) using SuperScript III reverse transcriptase (#18080044, Invitrogen) according to the manufacturer's protocol. The resulting cDNA was amplified in 30 PCR cycles with Phusion High-Fidelity DNA polymerase (#M0530, NEB) using primers 3 and 4 (primer 4: 5'- GGG TTA TGC TAG TTA TTG CTC AGC G). Purified DNA was digested with NdeI and XhoI restriction enzymes and ligated into the pMG-dB acceptor vector.

The resulting plasmid library was then employed in the next cycle, carried out as above, but in the presence of RNase A (10 µg/mL; #R4875, Sigma-Aldrich) instead of benzonase. The surviving variants were then subjected to a third cycle, with two modifications of the selection protocol. First, gel filtration was carried out on a HiPrep 16/60 Sephacryl-S400 (GE Healthcare) column, which has poorer resolution than the previously used column, but its higher exclusion volume allows efficient removal of larger aggregates. Second, nuclease treatment was extended to 4 hours and performed with a mixture of RNase A (10 µg/mL) and 2 vol% of an RNase cocktail enzyme mix from Thermo (#AM2288). From variants surviving the third selection cycle, 12 clones were picked, sequenced, and produced in E. coli. After affinity purification and SEC, the best variant in terms of yield, RNA packaging, minimal aggregation, and structural homogeneity—NC-4—was chosen for further analysis.

Production and purification of NC-1, NC-2, NC-3, and NC-4

All NCs were produced in E. coli BL21(DE3)-gold cells. Two-liter Erlenmeyer flasks containing 800 mL LB medium were inoculated with 8 mL overnight cultures and incubated at 37 °C and 200 rpm until the OD600 reached 0.5–0.7. Protein production was induced by adding IPTG to a final concentration of 0.5 mM. Cells were cultured at 25 °C for 18 hours and then harvested by centrifugation at 6,000 *g* and 15 °C for 20 min. The cell pellet from one 800-mL culture was resuspended in 20 mL LB medium, transferred and split into two 50-mL Falcon tubes. The medium used for transfer was removed by centrifugation at 4,000 *g* and 15 °C for 10 min, decanted, and aliquots of the cell pellet were frozen in liquid nitrogen and stored at −20 °C until purification. For purification, a cell pellet corresponding to 400 mL of culture volume was resuspended in 20 mL lysis buffer (50 mM sodium phosphate buffer at pH 7.4) containing 20 mM imidazole, and either 200 mM (NC-3), 500 mM (NC-1 and NC-2), or 1 M (NC-4) NaCl. The lysis buffer was supplemented with lysozyme (1 mg/mL). The mixture was incubated at room temperature for 20 min on an orbital shaker. After lysis by sonication (5 cycles of 1 min on, 1 min off, with amplitude = 80 and cycle = 60, UP200S sonicator, Hielscher Ultrasonics GmbH) and clearance by centrifugation at 8,500 *g* and 15 °C for 25 min, the supernatant was loaded onto 3 mL of Ni(II)-NTA agarose resin in a gravity flow column. After incubation for 10 min and washing with lysis buffer containing 20 mM imidazole, NCs were eluted with elution buffer (50 mM sodium phosphate buffer at pH 7.4, 500 mM imidazole) containing 200 (NC-3) or 500 mM (NC-1, NC-2, and NC-4) NaCl. The eluted fractions were concentrated and buffer-exchanged into storage buffer containing 200 mM (NC-3 and NC-4) or 500 mM (NC-1 and NC-2) NaCl using Amicon Ultra-15 centrifugal filter units (100 kDa MWCO, Merck Millipore). Protein capsids were further purified by SEC at room temperature using a Superose 6 increase 10/300 GL column equilibrated in storage buffer containing 200 (NC-3, NC-4) or 500 mM (NC-1, NC-2) NaCl. Purified fractions were pooled, concentrated, aliquoted, and either analyzed immediately, or frozen in liquid nitrogen and stored at –80 °C. Where stated, NC-4 was further purified by anion exchange chromatography at room temperature using a MonoQ 10/100 column (Pharmacia Biotech). The mobile phase consisted of storage buffer containing 200–1000 mM NaCl.

For NC-3 and NC-4, protein and RNA concentrations were measured by UV absorbance and deconvoluted using a previously reported protocol.[2] For NC-1 and NC-2, this calculation could not be applied, likely because scattering from aggregating particles skewed absorbance values. Extinction coefficients for proteins were calculated using the ExPASy ProtParam tool.[3] Wild-type AaLS was produced and purified as previously reported.[4]

Negative-stain transmission electron microscopy (TEM)

Negative-stain TEM was performed as reported previously.[5] Briefly, TEM grids (#01814-F, Ted Pella, Inc.) were negatively glow discharged at 15 mA for 45 s with a Pelco easiGlow Glow Discharge Cleaning System. After FPLC purification, grids were incubated with the capsid solution (10 μM monomer in storage buffer containing 200 mM NaCl) for 1 min, washed twice with doubly distilled water (ddH$_2$O), and once with TEM staining solution (2% wt/vol aqueous uranyl acetate, pH 4), after which the grids were incubated with staining solution for 10 s, dried, and imaged using a TFS Morgagni 268 microscope.

In vitro transcription of reference mRNAs

Reference messenger RNAs (mRNAs) for real-time quantitative PCR (RT-qPCR) were prepared by runoff in vitro transcription. DNA templates were prepared by PCR with primer 5 (primer 5: 5'- GCG AAA TTA ATA CGA CTC ACT AAT AG) and primer 6 (primer 6: 5'- CAA AAA ACC CCT CAA GAC CC) from plasmids pMG-dB-NC-1 to pMG-dB-NC-4 using the LongAmp Taq assay (#M0287, NEB). PCR-amplified templates were gel-purified using the DNA Clean & Concentrator-5 kit. In vitro transcription reactions were performed using T7 RNA polymerase (#EP0111, Thermo Scientific) according to the manufacturer's protocol. Template DNA was digested by RQ1 DNase and RNA was precipitated with isopropanol. RNA samples were purified twice by denaturing polyacrylamide electrophoresis (PAGE). Briefly, preparative urea PAGE gels (20 cm x 16 cm x 0.1 mm) were prepared in Tris/borate/EDTA (TBE) buffer supplemented with 8 M urea and 5% polyacrylamide. Polymerization was initiated using TEMED (8 μL per 10 mL gel solution) and APS (10% in water, 90 μL per 10 mL gel solution). RNA bands were visualized by UV shadowing and excised with a scalpel. The gel pieces were crushed with a pipet tip and the RNA was extracted in water containing 0.3 M NaCl overnight at room temperature. The next day, the RNA was purified by ethanol precipitation and dissolved in water. RNA quality and purity were assessed by measuring A260/A280 and A260/A230 ratios (for pure RNA, both ratios are ≥2.0) and by analytical PAGE gels. RNA concentrations were measured using the Qubit RNA HS assay (#Q32852, Invitrogen).

Extraction of nucleocapsid RNA and RT-qPCR

RNA was extracted from 100-µL or 200-µL aliquots of purified NCs containing a total amount of 5–10 µg RNA using the RNeasy Mini kit (#74104, QIAGEN) following the manufacturer's instructions. RNA standards were prepared by in vitro transcription as described above. After ensuring that RNA samples were free of contaminants by absorbance, concentrations from extracted RNAs and in vitro-transcribed standards were measured with the Qubit RNA HS Assay. cDNA of the capsid's genome was prepared by reverse transcription with primer 7 (5'- CCA AGG GGT TAT GCT AGT TAT TGC TCA GC) and SuperScript III reverse transcriptase (#18080044, Invitrogen) according to the manufacturer's protocol. After the reverse transcription reaction, RNase H (#18021014, Invitrogen) was added to digest RNA transcripts. Immediately following the reverse transcription reaction, dilutions of the cDNA were mixed with KOD SYBR qPCR Master Mix (#QKD-201, TOYOBO), primers 8 (5'- TGT GAG CGG ATA ACA ATT CCC CTC) and 9 (5'- GGG TTA TGC TAG TTA TTG CTC AGC G), and ROX reference dye according to the manufacturer's protocol. cDNA was amplified in 40 PCR cycles on a StepOnePlus thermocycler (Applied Biosystems) employing the thermocycler-specific PCR conditions provided in the qPCR mix manual. Absolute amounts of full-length genome were determined using standard curves prepared with cDNA originating from highly pure in vitro-transribed reference RNAs. The full RT-qPCR experiments to quantify the fraction of full-length mRNA in the total isolated RNA were repeated in two separate laboratories (by Angela Steinauer at ETH Zurich and by Naohiro Terasaka at the University of Tokyo) to ensure reproducibility.

Long-read sequencing

Nanopore sequencing was performed as described previously.[1] Oxford Nanopore Technology relies on polyadenylated RNAs. Therefore, the extracted NC RNAs were polyadenylated using E. coli poly(A) polymerase (#M0276, NEB) and purified using the RNA Clean & Concentrator-5 kit (#R1015, Zymo Research). cDNA libraries were prepared with the Direct cDNA Sequencing Kit (#SQK-DCS109, Oxford Nanopore Technologies) and Native Barcoding Kit 1D (#EXP-NBD104, Oxford Nanopore Technologies) following the manufacturer's protocols. Sequencing was carried out in a flow cell (#FLO-MIN106) using the 72-h 1D protocol. Base calling and de-multiplexing were performed using Oxford Nanopore Technology's Guppy Basecalling Software (version 3.2.10+aabd4ec). Adapter sequences of demultiplexed reads were removed using Porechop (version v0.2.4, https://github.com/rrwick/Porechop). Reads were mapped to the plasmid reference genome and to the E. coli genome (RefSeq: NC_000913.3) using Minimap2 (version 2.17 (r941), https://github.com/lh3/minimap2). Index reference files containing the pMG plasmid genomes and the E. coli genome were prepared using samtools (version 1.10, https://github.com/samtools/). Index reference files and mapped reads were imported into CLC Genomics Workbench (version 12.0, QIAGEN Bioinformatics). Alignments were sorted and the read sequences and lengths corresponding to the most abundant gene classes were extracted using samtools. For each gene, we calculated the sum of all gene-specific base pairs and compared it to the sum of all recorded base pairs.

Nuclease challenge assay

Aliquots of nucleocapsids containing a total amount of 5–10 μg RNA were treated in 50 mM sodium phosphate buffer at pH 7.4, 200 mM NaCl, 5 mM EDTA, 5 mM $MgCl_2$ either lacking nuclease or supplemented with benzonase (2.5 U/μL; 101654, Merck Millipore) or RNase A (10 μg/mL; #R4875, Sigma-Aldrich). The concentration of RNA and protein was held constant at 80 ng RNA/μL, which corresponds to about ~5 μg protein/μL. Aliquots were challenged with the respective nucleases at 37 °C for the indicated time periods after which samples were frozen in liquid nitrogen and stored at –80 °C. For analysis, RNA was extracted from the capsid as previously described.[6] A nuclease-treated NC solution (100 μL) was mixed with TRIzol (500 μL), vortexed for 3–5 s, and left on ice for 10 min. Then, chloroform (100 μL) was added, the samples were vortexed again, and the two phases were separated by centrifugation at 15,000 *g* for 20 min. The upper, aqueous layer (~300 μL) was carefully transferred to a clean Eppendorf tube and mixed with an equal volume of 20% ethanol in nuclease-free water. This extraction step was essential to remove nuclease contamination. Subsequently, the solution was transferred to an RNeasy Mini spin column, and RNA purified according to the manufacturer's protocol.

RNA stability was visualized on denaturing PAGE gels. Analytical urea PAGE gels (8.3 cm x 7.3 cm x 0.1 mm) were prepared in TBE buffer supplemented with 8 M urea and 8% polyacrylamide. Polymerization was initiated using TEMED (8 μL per 10 mL gel solution) and ammonium persulfate (APS) (10% in water, 90 μL per 10 mL gel solution). Gels were loaded with equal volumes of extracted RNA. The NC genome was selectively visualized using the fluorogenic dye DFHBI-1T (#446461, United States Biological), which fluoresces upon binding to the Broccoli aptamer that is part of the BoxBr tags.[7] Total RNA was visualized using GelRed (#41002, Biotium).

Cryo-electron microscopy: data collection and image processing

Freshly purified NCs were concentrated in storage buffer. NC-1 and NC-2 eluted as two major peaks from the SEC column, both of which were pooled for analysis. These variants were concentrated to a 280-nm absorbance of 20–30, as the protein concentration could not be estimated accurately due to the high absorbance ratio of 260/280 nm mentioned above. NC-3 and NC-4 were concentrated to 4–5 mg/mL. Copper-supported holey carbon grids (R2/2 Cu 400, Quantifoil) were negatively glow discharged at 15 mA for 15 s with a Pelco easiGlow Glow Discharge Cleaning System. Then, 3.5 μL of sample were applied and blotted with a vitrobot (FEI) for 12 to 14 s at 25 blot strength, 100% humidity, and 22 °C. Grids were plunged into liquid ethane and stored in liquid nitrogen.

Initial screening for all capsids and data collection for NC-3 were performed with a TFS Tecnai F20 equipped with a Falcon II direct electron detector (FEI). Movies of 7 frames were collected at a total dose of 40 electrons per Å2 and a magnification of 62,000x (1.8 Å pixel size). Defocus ranged from –1.8 to –3.3 μm. NC-1, NC-2, and NC-4 data collection was performed on a Titan Krios equipped with a Falcon III direct electron detector (FEI). Movies of 40 frames were collected at a dose of 60 electrons per Å2 and a magnification of 130,000x (1.1 Å pixel size). NC-4 was collected in electron counting, NC-1 and NC-2 in integration mode. Defocus ranged from –0.8 to –2.7 μm.

All single-particle reconstructions were performed in Relion 3.0 (ref. 8). Motion correction was performed with MotionCor2 (ref. 9) implemented in Relion, contrast transfer function (CTF) estimation with GCTF.[10] Good micrographs were selected based on metadata values and manual inspection. For NC-1, NC-2, and NC-3, early classifications were performed with CTF ignored up to the first peak to avoid grouping into a few, featureless classes.

Reconstruction of NC-1 and NC-2 structures was complicated by heterogeneity and aggregation. 2D classification was performed with multiple different mask sizes in order to obtain classes with distinct features for differently sized species. These classes were then used for the generation of initial 3D models of the tetrahedrally symmetric 120-mer (NC-1) and 180-mers (NC-1 and NC-2). For the final reconstruction though, as shown in Supplementary Figures 4 and 5, 2D classification was performed with a single large mask and size differences mainly separated subsequently in 3D classification, as this procedure led to higher particle numbers and consequently better resolution. We tried to reconstruct additional 3D structures from the heterogenous particles, but no other reasonable models could be obtained. Single- or multi-reference 3D classification based on known AaLS-derived structures, such as the 240-subunit capsid, or hollow spheres were not successful. The inability to extract further structures from the samples likely reflects substantial aggregation, low particle numbers of individual capsid architectures, shape irregularities, and lower symmetry.

In contrast to NC-1 and NC-2 particles, NC-3 and NC-4, were better behaved and more homogeneous, making data elaboration according to standard procedures[8] fairly straightforward. Good 2D classes were used to generate initial models with imposed icosahedral symmetry. The best classes from 3D classification, masked around the capsid shells, were further refined. Further processing steps are described in Supplementary Figure 6.

Model building and refinement were performed in Coot 0.8.9.2,[11] Phenix 1.18,[12] and Pymol 2.0. Electron density maps from 3D refinement, postprocessing in Relion, and autosharpening in Phenix were used during model building. NC-1, NC-2, and NC-4 models were based on a crystal structure of the wild-type lumazine synthase (PDB-ID: 1hqk).

Atomic models were initially built into the asymmetric units and refined. After symmetry expansion, the full capsids were refined with non-crystallographic symmetry constraints to reflect the symmetry imposed during reconstruction. Experimental data versus model geometry were weighted in Phenix to optimize both electron density fit and geometry. While the core fold of the protomers in the tetrahedrally symmetric capsids were well resolved, the maps for the segment encompassing residues 66–81 displayed lower local resolution in subunits where this area is exposed towards the capsid openings and not in contact with neighboring protomers. These segments were built by repositioning the known structural elements of the wild-type protein as rigid groups and remodeling the flanking linkers according to visible density and chemical constraints, although multiple alternative conformations may exist. The pseudo-atomic NC-3 model is based on the structure of NC-4 with reversion of the mutations and an additional cycle of refinement to satisfy geometric and steric constraints. More information on data collection and model building is found in Table S1.

XRF data collection and analysis

XRF experiments were performed as described[13] and analyzed using QuShape[14] modified to incorporate sample replicate comparisons. In vitro-transcribed and NC-packaged RNAs were exposed in triplicate to X-ray pulses of 25 or 50 ms at the National Synchrotron Light Source II, beamline 17-BM XFP at Brookhaven National Laboratory (Upton, NY). Packaged RNA was subsequently extracted from the protein shell by standard techniques.[13]

Nucleotide modification propensity (reactivity) is directly related to residue mobility, and thus reflects base pairing and inter-molecular contacts. Reactivity was quantitated post-exposure by capillary electrophoresis sequencing using three dye-labeled primers (primer 10: 5’- CCA AGG GGT TAT GCT AGT TAT TGC TCA GC; primer 11: ATG CTA CGA TAC CGA AAC GAA GGC; primer 12: 5’- CTC GAT AGC CTG TTC CAA GGT G) that cover ~90% of the genome, including the coding region of the structural protein. Pairwise Pearson correlation coefficients (PCC) for normalized replicates gave best correlations overall at 50 ms exposure (Table S3), and the respective data were therefore further analyzed. XRF footprints showing normalized reactivities for each nucleotide were generated for both the free and packaged state using established protocols.[15,16]

To find potential packaging signals (PSs), we looked for sequences similar to the UGxAxAA motif (x, any nucleotide) and the UGxA submotif, which are known to bind the λN+ peptide.[17] Eighteen occurrences of URxRxRR (R, any purine) and 21 of URxRxxx were identified in the mRNA transcripts of both NC-3 and NC-4 (Table S2). Contact with the RNA-binding peptide would result in lowered reactivity. Of the 39 identified sites, only 13 (highlighted in grey in Table S2) displayed such low reactivity levels (green and black colored nts in Table S2), including the two copies of the BoxB sequence (BB1 & BB2).

XRF reactivity levels were used as constraints to weight RNA secondary structure predictions. We used a modification of the RNA folding algorithm S-fold that includes such data via a scaling factor ($m$) and an offset ($b$) to generate a statistical sample of secondary structures from the Boltzmann ensemble of RNA secondary structures.[15] Typically, the ($m$,$b$) combination that best represents a known secondary structure element within a probed RNA is identified, and that combination is used to predict the overall secondary structure.[16] Because the reported stem-loop of the BoxBr tag contains C-G base pairs that stabilize the stem,[18] it occurs with high probability in the ensembles for many (*m*,*b*) combinations, and it is therefore insufficiently discriminatory to identify a unique (*m*,*b*) combination. We therefore computed 1000 statistical (Boltzmann-weighted) sample folds for all 1116 (*m*,*b*) combinations for *m* values between 0 and 7 and *b* values between 0 and -6, in increments of 0.2. Computing multiple folds per (*m*,*b*) combination takes into consideration that large RNAs occur as ensembles of secondary structures with comparable folding free energies. For any of the sites to act as packaging signals, they must be presented with sufficient frequency in the ensemble. In order to identify trends, the maximum, minimum and average frequency of stem-loops overlapping with the identified motifs were computed over all sample folds and all (*m*,*b*) combinations tested, both for the in vitro transcript and packaged RNA. Of the 13 identified sites, only seven (BB1, BB2, and PS1–5; Table S2) appear as part of a loop in either NC-3 or NC-4 with significant frequency (>50% of the folds in >50 of the (*m*,*b*) combinations).

For the calculation of representative folds of the packaged NC-3 and NC-4 mRNA, (*m*,*b*) values were chosen taking into account contributions from all PSs which were preferentially displayed in the packaged over free transcripts via their cumulative normalized frequencies of occurrence (Supplementary Figures 8,9). Two local probability maxima were identified for both packaged mRNAs, and structures computed for both. Maximum ladder distances for these folds (Supplementary Figures 8,9) show that evolution did not select for genome compactness, likely because the mRNA is considerably smaller than the packaging capacity of the evolved T=4 capsids.

#### Supplementary Figures and Tables

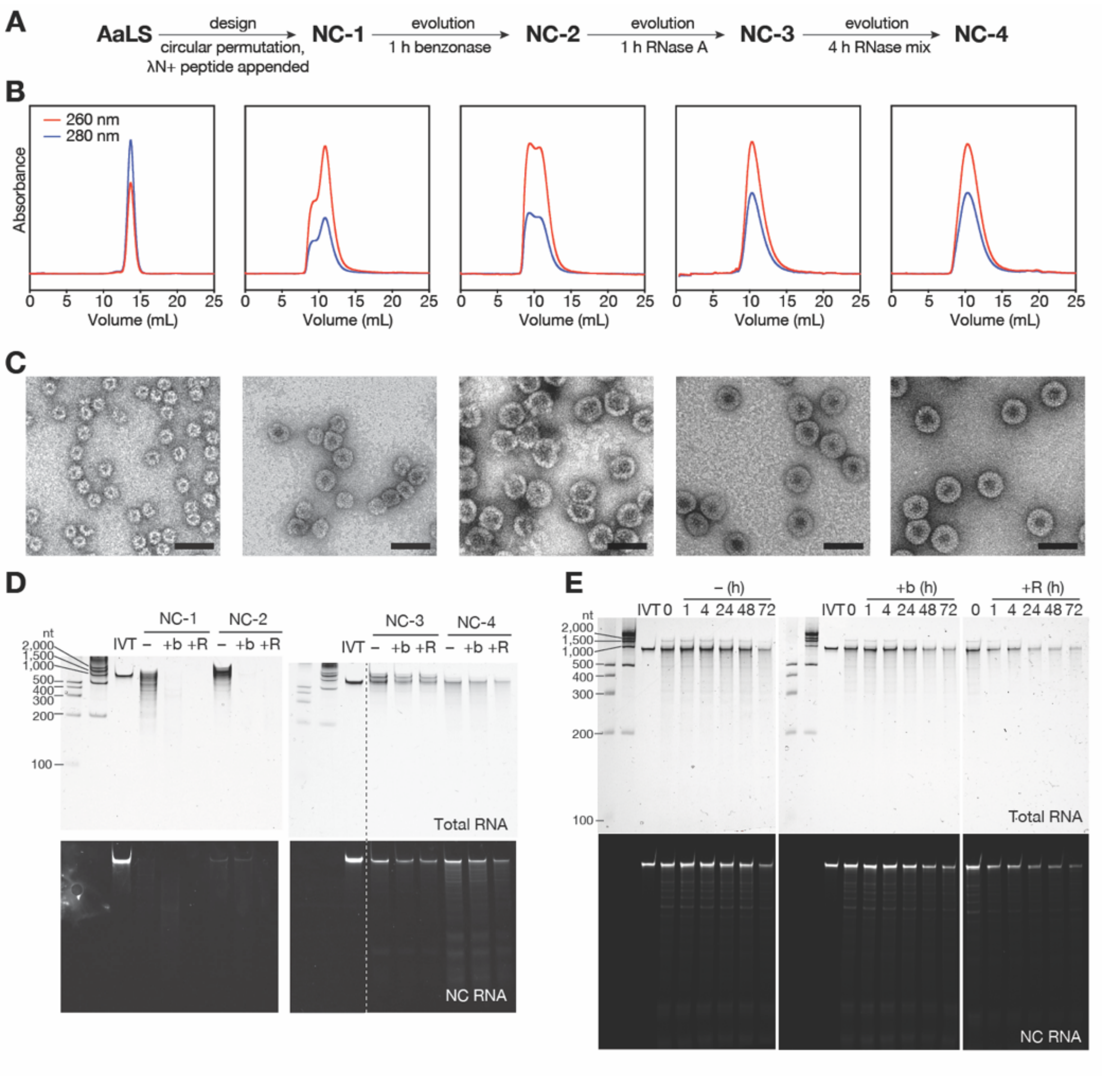


**Supplementary Figure 1. Nucleocapsid design and evolution.**

(A) NC-1 was generated by circular permutation of AaLS and addition of the λN+ peptide to the new N-terminus.[1] Directed evolution over three generations with increasingly stringent nuclease challenge in each step yielded NC-4. As explained in the Materials and Methods section, the previously described nucleocapsids were renamed to clarify their evolutionary relationships. (B) Size-exclusion chromatograms of purified, re-injected nucleocapsids (column: Superose 6 increase 10/300 GL). (C) Transmission electron micrographs of purified capsids. Scale bar: 50 nm. (D) Capsid stability towards nucleases: Purified NCs were incubated without nuclease (–), or treated with benzonase (+b), or RNase A (+R) for 1 hour at 37 °C. RNA was extracted and equal volumes were loaded onto a denaturing PAGE (8%) gel. Total RNA was stained with GelRed, nucleocapsid mRNA (NC RNA) was visualized with DFHBI-1T, a small molecule that fluoresces upon binding to the broccoli aptamer present in the 5'- and 3'-untranslated regions of the capsid mRNA. IVT = in vitro-transcribed reference mRNA. The dashed line indicates two non-concurrent portions of the same gel image. (E) Purified NC-4 was treated with nucleases as in (D) for the indicated number of hours and analyzed on a denaturing PAGE (5%) gel.

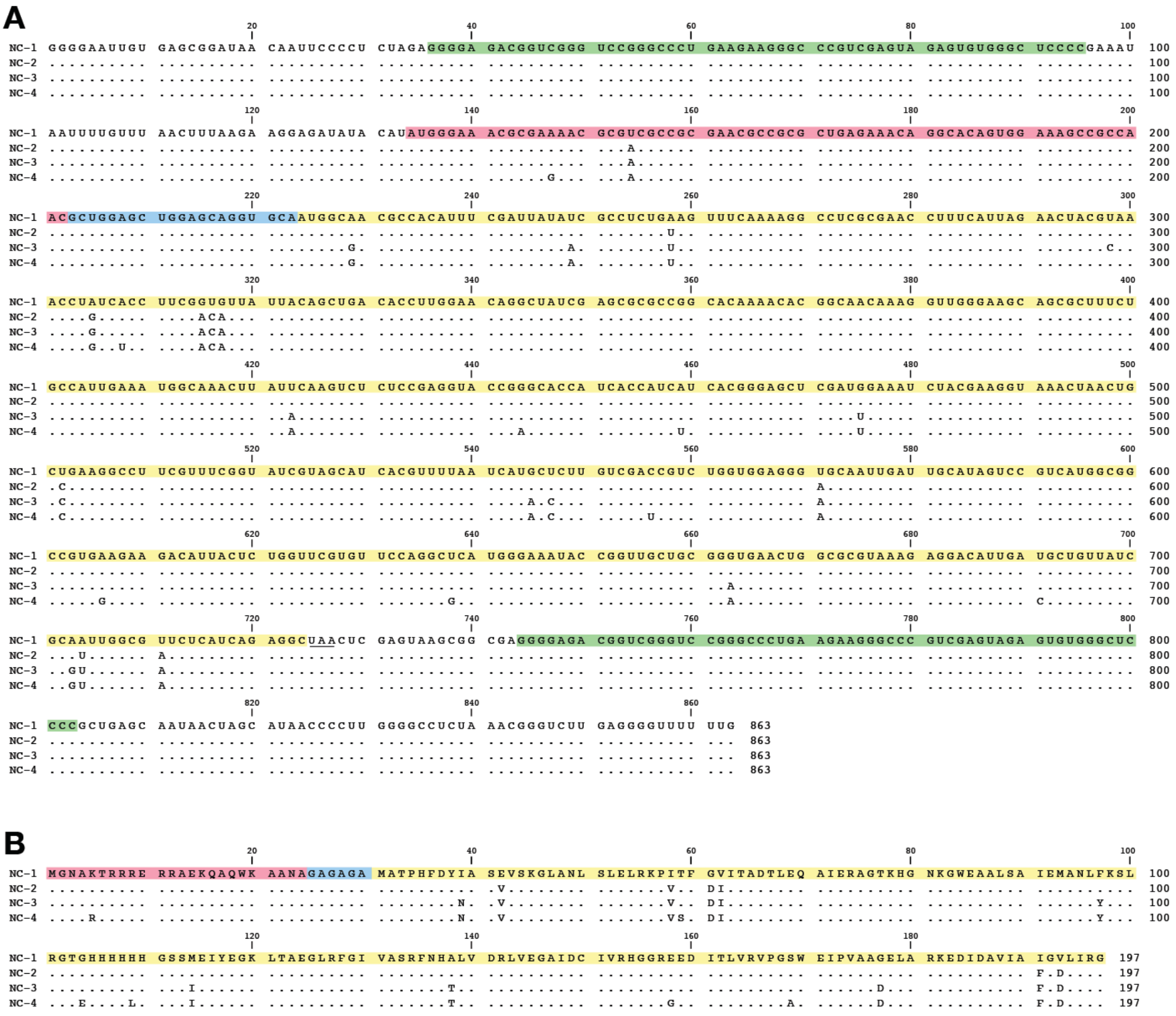


**Supplementary Figure 2. Sequence alignment of NC-1 to NC-4.**

(A) mRNA and (B) protein sequences of NC-1 to NC-4 (green = BoxBr tags, magenta = λN+ peptide, blue = (GlyAla)-linker, yellow = cpAaLS).

**A**

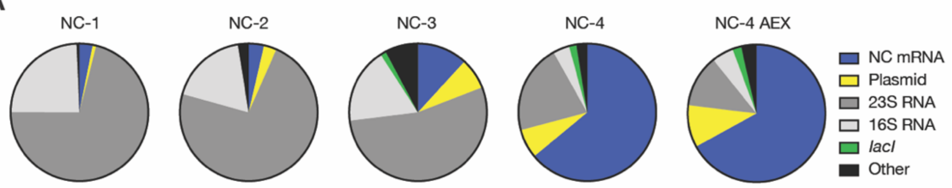


**B**

| NC-1 | | NC-2 | | NC-3 | | NC-4 | | NC-4 AEX | |
|---|---|---|---|---|---|---|---|---|---|
| Gene | % bp | Gene | % bp | Gene | % bp | Gene | % bp | Gene | % bp |
| NC-1 | 3.100 | NC-2 | 3.600 | NC-3 | 11.800 | NC-4 | 64.000 | NC-4 | 67.000 |
| Plasmid | 0.830 | Plasmid | 3.008 | Plasmid | 7.320 | Plasmid | 6.864 | Plasmid | 9.945 |
| 23S rRNA | 71.128 | 23S rRNA | 72.704 | 23S rRNA | 53.952 | 23S rRNA | 21.015 | 23S rRNA | 12.293 |
| 16S rRNA | 24.250 | 16S rRNA | 18.166 | 16S rRNA | 17.829 | 16S rRNA | 3.851 | 16S rRNA | 5.103 |
| *lacI** | 0.090 | *lacI** | 0.269 | *lacI** | 1.335 | *lacI** | 1.757 | *lacI** | 2.078 |
| Other: | | Other: | | Other: | | Other: | | Other: | |
| *rnpB* | 0.003 | *rnpB* | 0.004 | *gltA* | 0.199 | *gltA* | 0.278 | *gltA* | 0.306 |
| *ssrA* | 0.002 | *ssrA* | 0.004 | *rnpB* | 0.093 | *acnB* | 0.149 | *acnB* | 0.227 |
| Other genes† | 0.597 | *gltA* | 0.004 | *gpmA* | 0.062 | *gpmA* | 0.077 | *idc* | 0.096 |
| | | *acnB* | 0.004 | *ssrA* | 0.039 | *serS* | 0.048 | *rnpB* | 0.095 |
| | | *idc* | 0.002 | *acnB* | 0.035 | *rnpB* | 0.039 | *gpmA* | 0.084 |
| | | *serS* | 0.002 | *dadA* | 0.017 | *idc* | 0.039 | *aspC* | 0.054 |
| | | *acpP* | 0.001 | *acpP* | 0.012 | *dadA* | 0.030 | *serS* | 0.045 |
| | | *dadA* | 0.001 | *aspC* | 0.011 | *ychF* | 0.025 | *dadA* | 0.044 |
| | | *gpmA* | 0.001 | *idc* | 0.008 | *aspC* | 0.024 | *acpP* | 0.030 |
| | | Other genes† | 2.230 | *ychF* | 0.005 | *acpP* | 0.010 | *ychF* | 0.021 |
| | | | | *serS* | 0.003 | *ssrA* | 0.003 | *ssrA* | 0.010 |
| | | | | Other genes† | 7.281 | Other genes† | 1.792 | Other genes† | 2.569 |
| Total | 100.000 | Total | 100.000 | Total | 100.000 | Total | 100.000 | Total | 100.000 |
| #Reads | 104,325 | #Reads | 81,784 | #Reads | 48,669 | #Reads | 92,166 | #Reads | 68,253 |

*The *lacI* gene is present in both the plasmid and the bacterial genome.
†Sum of all remaining bps aligned with the *E. coli* genome present at <0.001%.

**C**

**D**

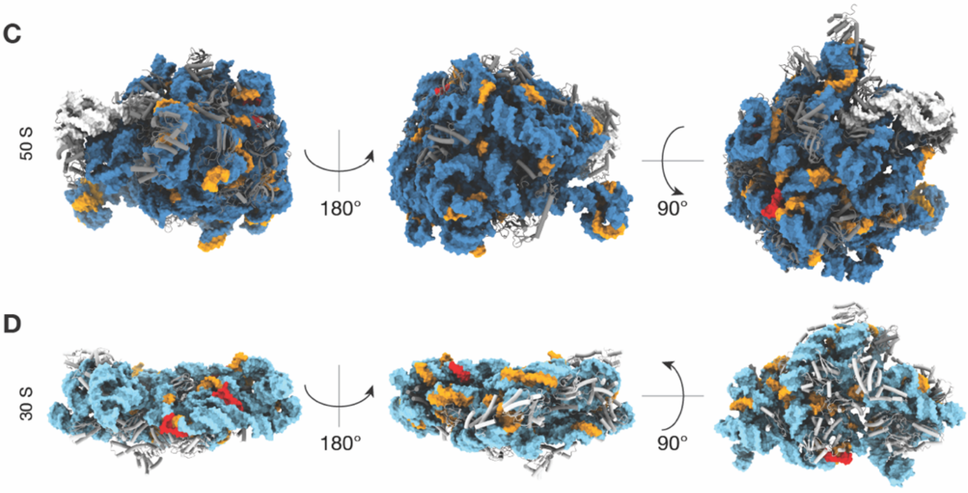


**Supplementary Figure 3. RNA cargo identified by long-read sequencing.**

(A) Pie chart representations of main gene classes identified by nanopore sequencing.[19] (B) Relative fraction of identified gene classes in percent calculated by adding gene-specific base pairs (bps) and comparing them to the sum of all recorded bps. (C and D) UGxAxAA (red) and URxRxRR (orange) motifs are mapped onto the 50S (C) and 30S (D) subunits of the E. coli ribosome (PDB: 5h5u). The 23S rRNA is colored in dark blue, the 16S rRNA in light blue, accessory proteins are shown as cartoons. The ubiquity and compactness of ribosomes, together with interactions between some of the exposed BoxB-like RNA motifs with the λN+ peptide, may explain competitive encapsidation of the ribosome by the nucleocapsids.

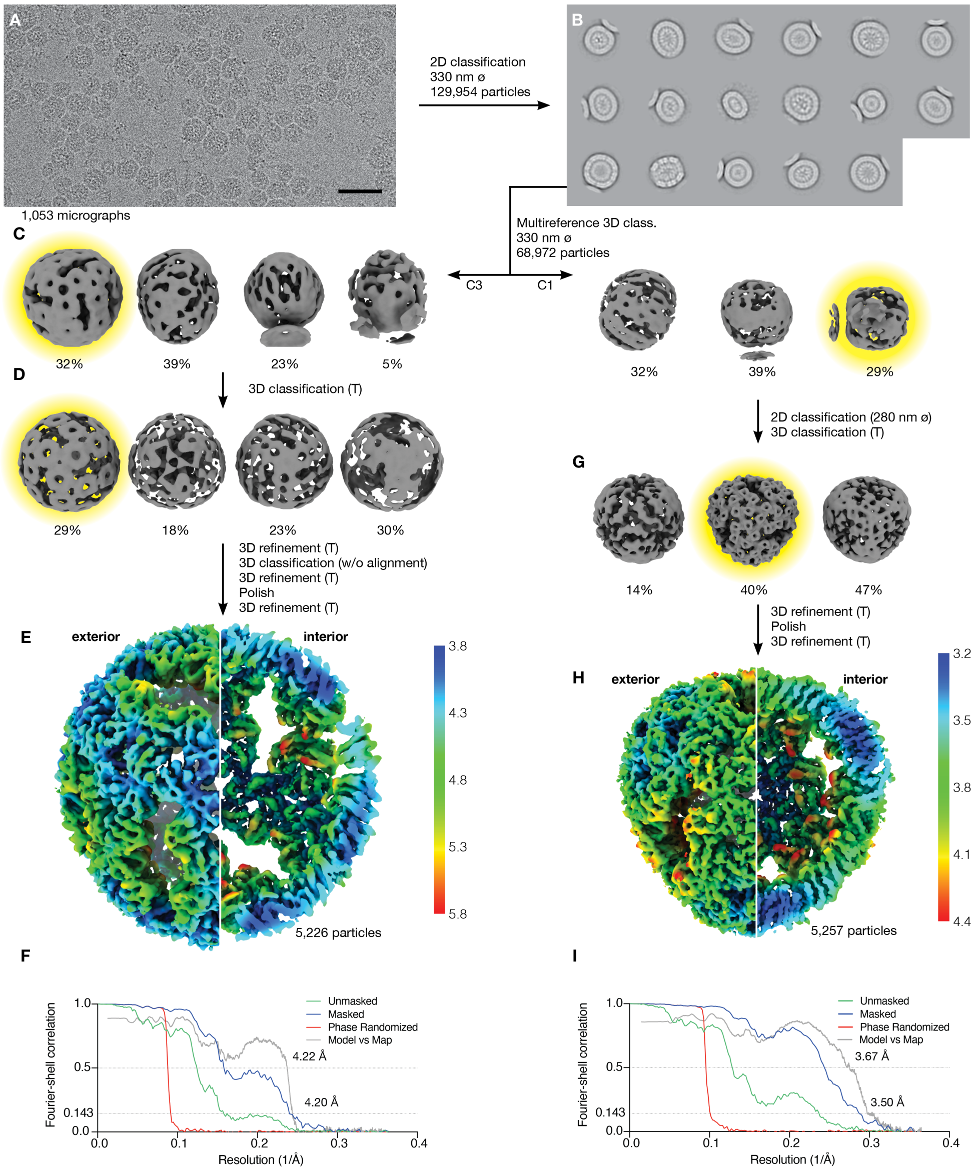


**Supplementary Figure 4. Single particle reconstruction of NC-1 structures.**

(A) 1,053 movies were analyzed for reconstruction of NC-1 (scale bar: 50 nm). (B) From these, 129,954 particles were picked and classified in 2D. (C) Size differences of the heterogeneous particles were initially classified in 3D with multiple references and lower symmetry (C1 and C3). References included the NC-4 structure (Supplementary Figure 6). (D and G), Particles were further classified with tetrahedral symmetry, using initial models generated beforehand by 2D classification with tight masks and then imposition of tetrahedral symmetry on classes showing distinct features. As indicated, further 2/3D classifications of classes highlighted in yellow, polishing, and refinement with imposed tetrahedral symmetry led to the final structures. (E and H) Refined maps, colored by local resolution, of the 180-mer (5,226 particles, 4.20 Å) (E) and the 120-mer (5,257 particles, 3.50 Å) (H). (F and I), Gold-standard Fourier-shell correlation curves for the 180-mer (F) and 120-mer (I).

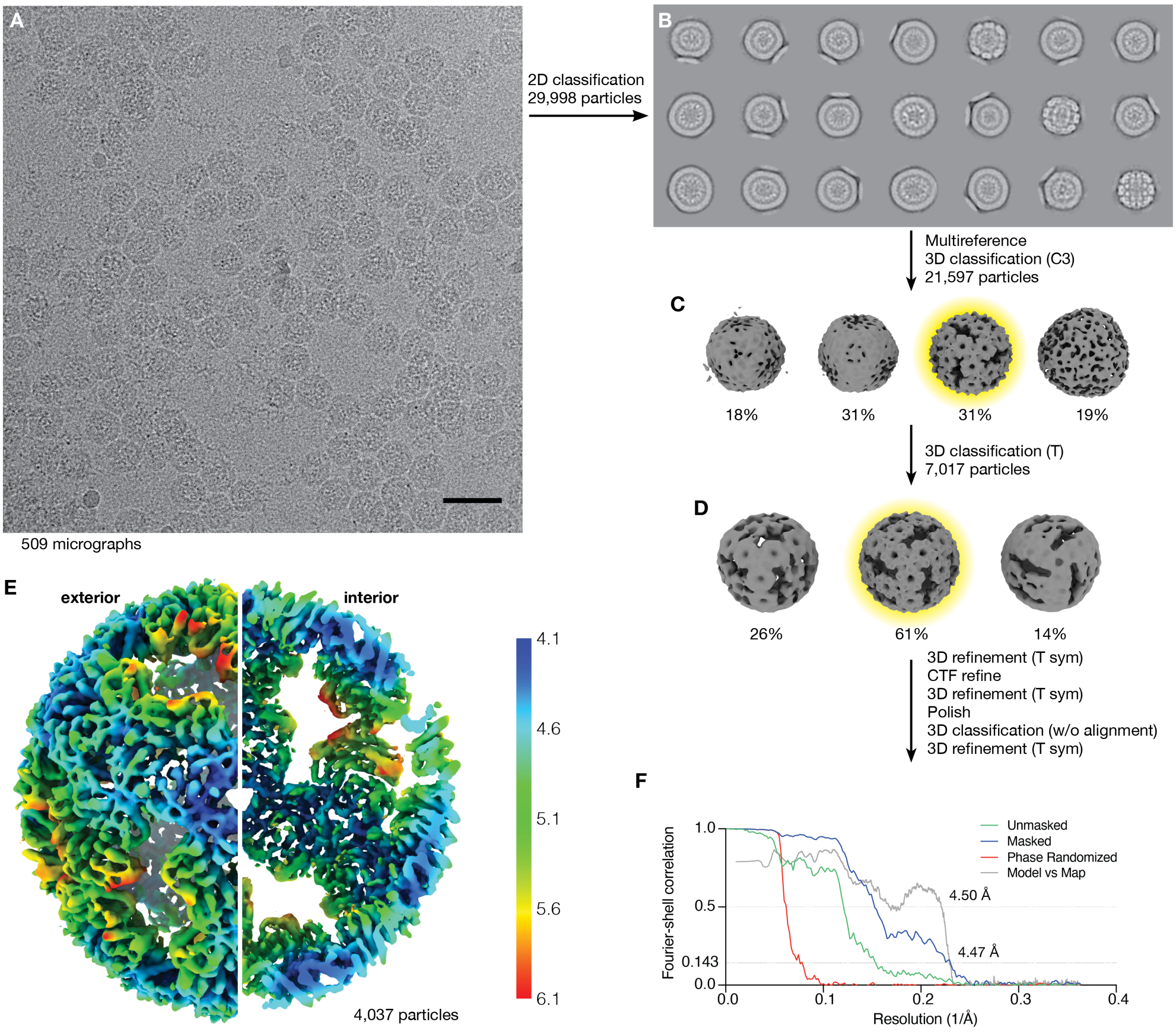


**Supplementary Figure 5. Single particle reconstruction of NC-2.**

(A) 509 movies were used to analyze NC-2 (scale bar: 50 nm). (B) 29,998 particles were classified in 2D. (C) Non-junk particles were further classified in 3D with multiple references (including the structure of NC-4, Supplementary Figure 6) with C3-symmetry. (D) Further 3D classification (with tetrahedral symmetry) of classes highlighted in yellow, contrast transfer function (CTF) refinement, polishing, and refinement with tetrahedral symmetry of individual classes led to the final structures. (E) Refined map, colored by local resolution (4,037 particles, 4.47 Å). (F) Gold-standard Fourier-Shell correlation curves.

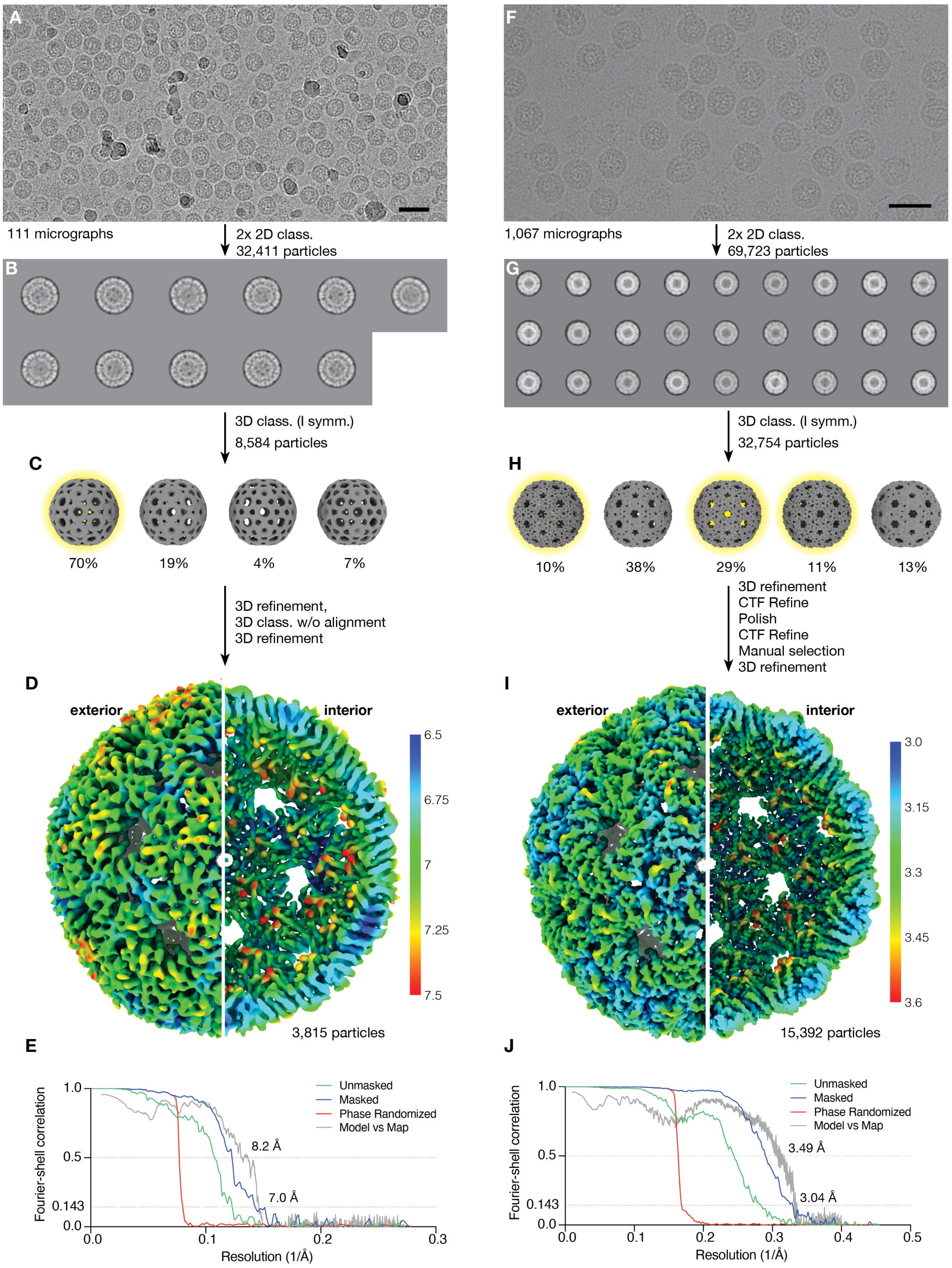


**Supplementary Figure 6. Single particle reconstruction of NC-3 (A–E) and NC-4 (F–J).**

(A and F) Sample movies from NC-3 (A) and NC-4 (F) micrographs (scale bar: 50 nm). (B and G) Particles were classified in 2D, and symmetric classes with clear features processed further. (C and H) Initial model generation and 3D classification were performed with imposition of icosahedral symmetry. For NC-4, multiple highly similar 3D classes (highlighted in yellow) were pooled. Particles were further processed as indicated. (D) Postprocessed map of NC-3 (3,815 particles, 7.0 Å). (I) Refined map of NC-4 (15,392 particles, 3.04 Å). (E and J) Gold-standard Fourier-Shell correlation curves.

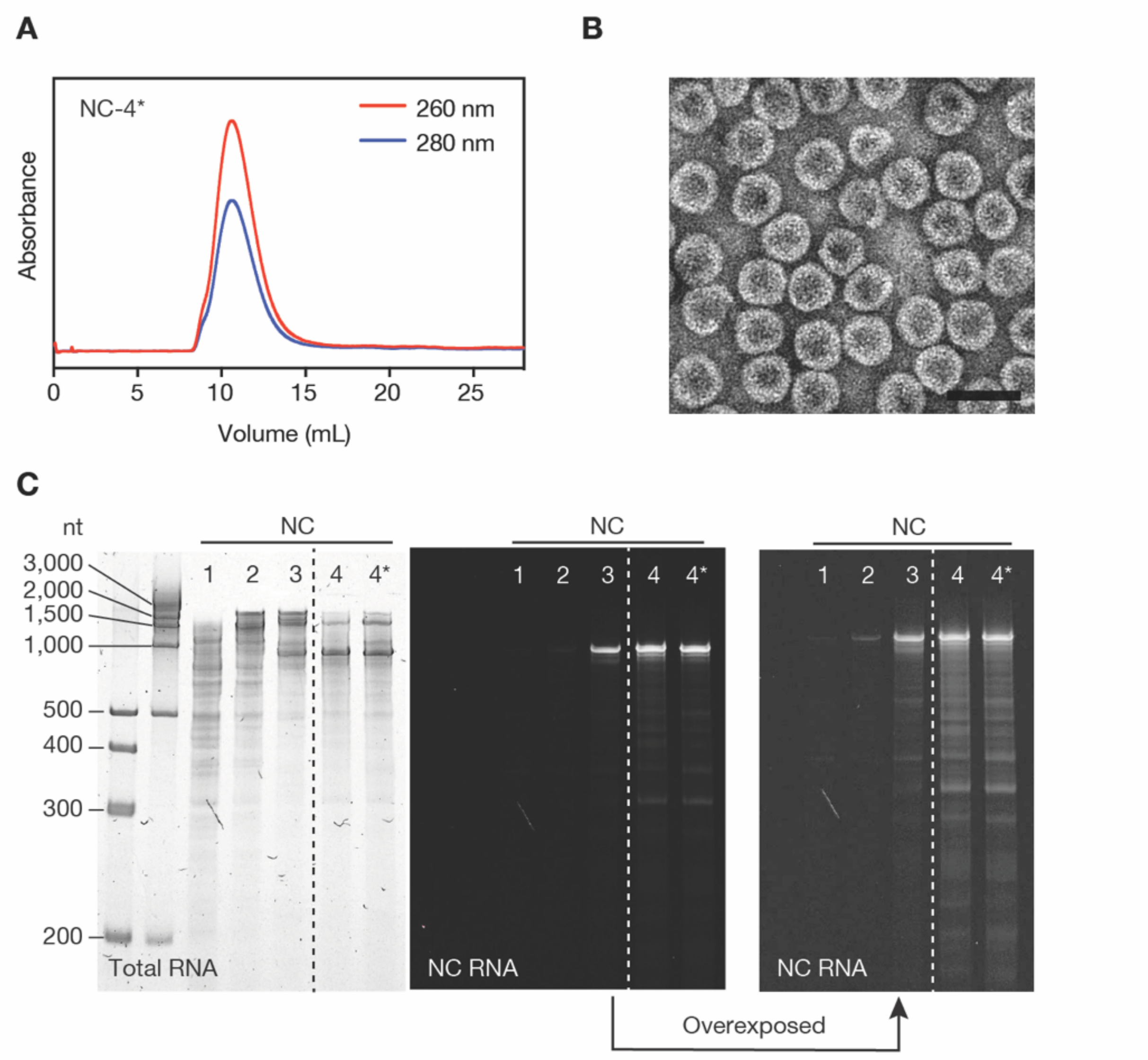


**Supplementary Figure 7. Reversion of the K5R mutation in NC-4.**

(A) Size-exclusion chromatograms of purified, re-injected nucleocapsid (column: Superose 6 increase 10/300 GL). (B) Transmission electron micrograph of purified R5K NC-4 (NC-4*). Scale bar: 50 nm. (C) RNA was extracted from each nucleocapsid generation and equal amounts were loaded onto a denaturing PAGE (5%) gel. Total RNA was stained with GelRed, NC RNA was visualized with DFHBI-1T. The DFHBI-1T-stained gel on the right was overexposed to better visualize faint bands. The dashed line indicates two non-concurrent portions of the same gel image.

**A**

| | Transcript | | | Packaged | | |
|---|---|---|---|---|---|---|
| | Max | Min | Avg | Max | Min | Avg |
| **PS1** | 1000 | 0 | 120 | 1000 | 0 | 578 |
| **BB1** | 1000 | 90 | 963 | 1000 | 0 | 628 |
| **PS2** | 922 | 0 | 37 | 1000 | 0 | 526 |
| **PS3** | 687 | 0 | 41 | 981 | 0 | 135 |
| **PS4** | 1000 | 2 | 679 | 1000 | 2 | 867 |
| **PS5** | 721 | 0 | 209 | 729 | 0 | 216 |
| **BB2** | 1000 | 0 | 750 | 1000 | 0 | 516 |

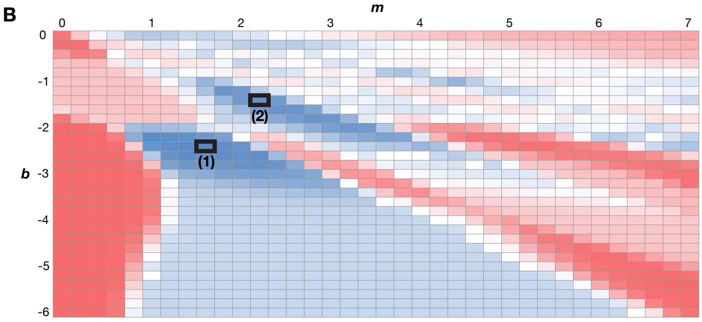


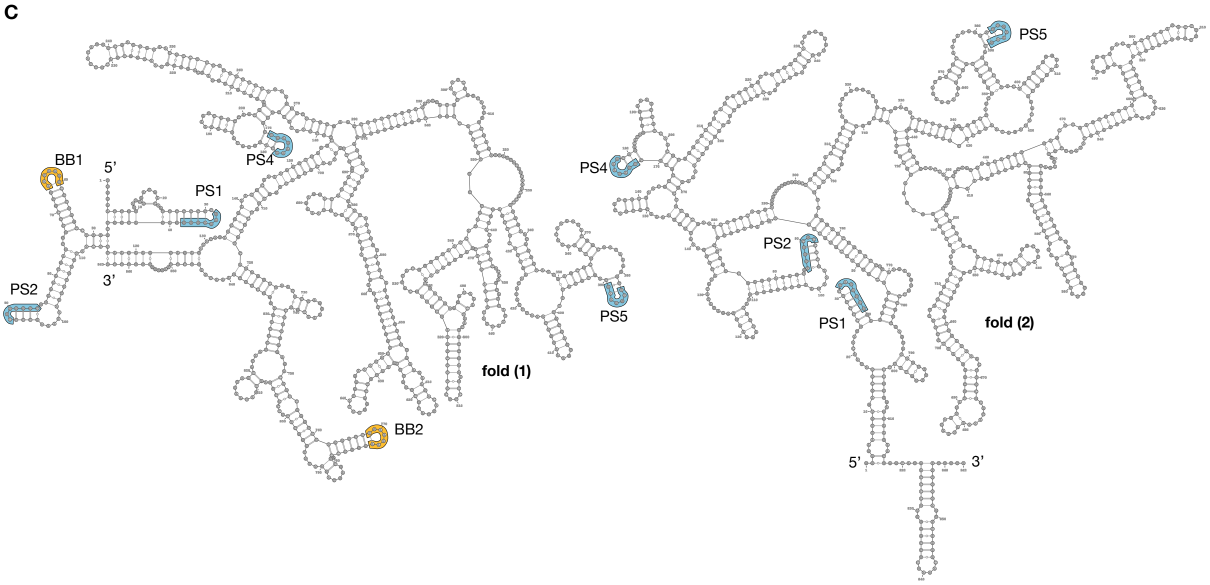


**Supplementary Figure 8. Secondary structure prediction based on XRF data for NC-3.**

(A) The minimum, maximum and average number of times a given SL occurs in an ensemble of 1000 sample folds, generated using a modified version of the S-fold algorithm that includes the XRF data via a scaling factor *m* and an offset *b*, was computed for each of 1116 (*m*,*b*) combinations (see panel B). The seven potential packaging signals listed occur in at least 50% of the 1000 sample folds for at least 50 of the 1116 (*m*,*b*) combinations. Stem-loops for which the average number increases for packaged mRNA compared with the free transcript, or vice versa, are highlighted; if the lower value is within 85%, it is highlighted in lighter shade. (B) Stem-loops with entries highlighted in packaged RNA in A are collectively optimized via a cost function given by the sum of the normalized (by their maximal number of occurrence) frequencies for these SLs in an ensemble of 1000 sample folds for each (*m*,*b*) combination. This identifies (*m*,*b*) values for which their occurrence is locally maximally aligned with the trend in the tables in A. The (*m*,*b*) combinations for which the cost function is maximal are: *m*=1.6, *b*=-2.4 (77.6%, labelled 1) and *m*=2.2, *b*=-1.4 (75.7%, labelled 2). (C) The predicted folds corresponding to these (*m*,*b*) combinations, represented as cartoons in Figure 4C, are shown with their full sequence. The maximum ladder distances are 98 for fold (1), and 102 for fold (2).

**A**

| | Transcript | | | Packaged | | |
|---|---|---|---|---|---|---|
| | Max | Min | Avg | Max | Min | Avg |
| **PS1** | 429 | 0 | 16 | 1000 | 0 | 460 |
| **BB1** | 1000 | 11 | 914 | 1000 | 794 | 990 |
| **PS2** | 900 | 0 | 80 | 1000 | 0 | 376 |
| **PS3** | 585 | 0 | 31 | 605 | 0 | 58 |
| **PS4** | 1000 | 1 | 691 | 1000 | 0 | 609 |
| **PS5** | 731 | 0 | 334 | 697 | 0 | 82 |
| **BB2** | 1000 | 0 | 844 | 1000 | 0 | 336 |

**B**

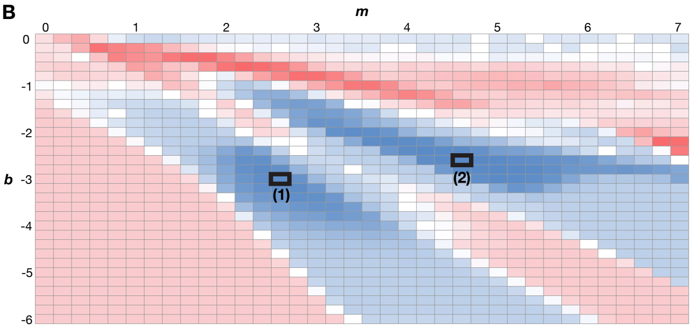


**C**

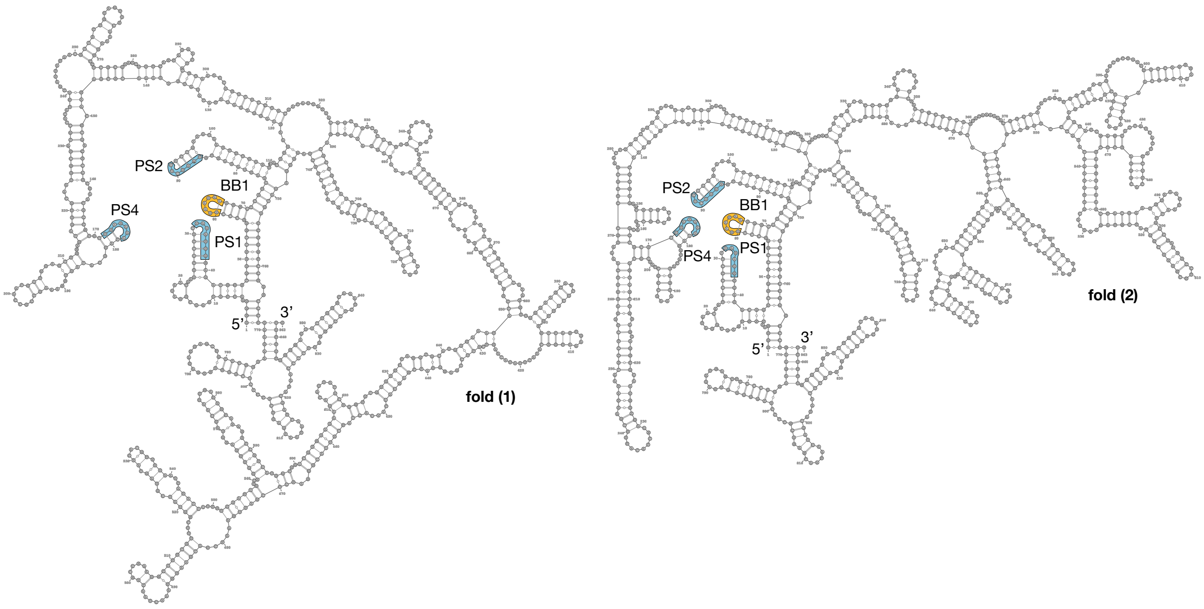


**Supplementary Figure 9. Secondary structure prediction based on XRF data for NC-4.**

(A) The minimum, maximum and average number of times a given SL occurs in an ensemble of 1000 sample folds, generated using a modified version of the S-fold algorithm that includes the XRF data via a scaling factor *m* and an offset *b*, was computed for each of 1116 (*m*,*b*) combinations (see panel B). The seven potential packaging signals listed occur in at least 50% of the 1000 sample folds for at least 50 of the 1116 (*m*,*b*) combinations. Stem-loops for which the average number increases for packaged mRNA compared with free transcript, or vice versa, are highlighted; if the lower value is within 85%, it is highlighted in lighter shade. (B) Stem-loops with entries highlighted in packaged RNA in A are collectively optimized via a cost function given by the sum of the normalized (by their maximal number of occurrence) frequencies for these SLs in an ensemble of 1000 sample folds for each (*m*,*b*) combination. This identifies (*m*,*b*) values for which their occurrence is locally maximally aligned with the trend in the tables in (A). The (*m*,*b*) combinations for which the cost function is maximal are: *m*=4.6, *b*=-2.6 (79.9%, labelled 1) and *m*=2.6, *b*=-3.0 (79.0%, labelled 2). (C) The predicted folds corresponding to these (*m*,*b*) combinations, represented as cartoons in Figure 4D, are shown with their full sequence. The maximum ladder distances are 125 for fold (1), and 105 for fold (2).

### Table S1. Cryo-EM data.

| | NC-1 120-mer | NC-1 180-mer | NC-2 | NC-3 | NC-4 |
|---|---|---|---|---|---|
| **EMDB map entry** | 11631 | 11632 | 11633 | 11634 | 11635 |
| **PDB coordinate entry** | 7A4F | 7A4G | 7A4H | 7A4I | 7A4J |
| ***Data Collection and reconstruction*** | | | | | |
| **Microscope model** | FEI Titan Krios | FEI Titan Krios | FEI Titan Krios | FEI Tecnai F20 | FEI Titan Krios |
| **Detector model** | Falcon III | Falcon III | Falcon III | Falcon II | Falcon III |
| **# of Micrographs collected** | 1481 | 1481 | 848 | 134 | 1080 |
| **Magnification** | 130 000x | 130 000x | 130 000x | 62 000x | 130 000x |
| **Voltage (kV)** | 300 | 300 | 300 | 200 | 300 |
| **Electron dose (e-/A2)** | 60 | 60 | 60 | 40 | 60 |
| **Pixel Size (Å)** | 1.1 | 1.1 | 1.1 | 1.8 | 1.1 |
| **Defocus range (µm)** | -0.8 to -2.6 | -0.8 to -2.6 | -0.8 to -2.6 | -1.8 to -3.3 | -0.8 to -2.6 |
| **Symmetry imposed** | T | T | T | I1 | I1 |
| **# of Micrographs used** | 1 053 | 1 053 | 509 | 111 | 1 067 |
| **Initial particle images** | 129 954 | 129 954 | 29 998 | 32 411 | 69 723 |
| **Final particle images** | 5 257 | 5 226 | 4 037 | 3815 | 15 392 |
| **Resolution (Å) (at FSC = 0.143** | 3.50 | 4.20 | 4.47 | 7.04 | 3.04 |
| **Map sharpening B-factor (Å2)** | -31 | -52 | -141 | -550 | -66 |
| ***Model building*** | | | | | |
| Starting model | 1hqk | 1hqk | 1hqk | 7a4j | 1hqk |
| **Composition** | | | | | |
| Chains | 120 | 180 | 180 | 240 | 240 |
| Atoms | 141432 | 205680 | 204396 | 290520 | 289560 |
| Protein residues | 18540 | 27120 | 26904 | 37860 | 37860 |
| Water | 0 | 0 | 0 | 0 | 0 |
| Ligands | 0 | 0 | 0 | 0 | 0 |
| **Bonds (RMSD)** | | | | | |
| Length (Å) (# > 4σ) | 0.002 (0) | 0.002 (0) | 0.002 (0) | 0.002 (0) | 0.002 (0) |
| Angles (°) (# > 4σ) | 0.430 (12) | 0.383 (34) | 0.396 (0) | 0.466 (189) | 0.402 (0) |
| **MolProbity score** | 1.83 | 1.46 | 1.67 | 1.86 | 1.58 |
| **Clash score** | 6.30 | 6.41 | 6.06 | 5.65 | 3.77 |
| **Ramachandran plot (%)** | | | | | |
| Outliers | 0 | 0 | 0 | 0 | 0 |
| Allowed | 2.36 | 2.27 | 1.81 | 2.57 | 1.82 |
| Favored | 97.64 | 97.73 | 98.19 | 97.43 | 98.18 |
| **Ramachandran plot Z-score** | | | | | |
| whole | 1.60 (0.06) | 2.89 (0.06) | 3.99 (0.05) | 1.36 (0.04) | 0.40 (0.04) |
| helix | 2.31 (0.05) | 2.64 (0.05) | 3.81 (0.04) | 2.05 (0.04) | 1.02 (0.04) |
| sheet | 0.45 (0.08) | 1.44 (0.07) | 2.32 (0.07) | 1.59 (0.07) | 1.39 (0.07) |
| loop | 0.02 (0.10) | 0.95 (0.09) | 0.73 (0.08) | 1.04 (0.05) | 1.28 (0.05) |
| **Rotamer outliers (%)** | 3.40 | 1.14 | 2.76 | 3.80 | 3.50 |
| **Cβ outliers (%)** | 0 | 0 | 0 | 0 | 0 |
| **Peptide plane (%)** | | | | | |
| Cis proline/general | 0.0/0.0 | 0.0/0.0 | 0.0/0.0 | 0.0/0.0 | 0.0/0.0 |
| Twisted proline/general | 0.0/0.0 | 0.0/0.0 | 0.0/0.0 | 0.0/0.0 | 0.0/0.0 |
| **CaBLAM outliers (%)** | 0.07 | 0.93 | 1.1 | 1.14 | 0.98 |
| **ADP (B-factors)** | | | | | |
| Iso/Aniso (#) | 141432/0 | 205680/0 | 204396/0 | 290520/0 | 289560/0 |
| min/max/mean | 45.97/167.52/91.60 | 14.53/153.78/58.76 | 34.68/253.39/95.22 | 96.48/410.28/208.34 | 41.87/138.99/82.97 |
| **Occupancy** | | | | | |
| Mean | 1 | 1 | 1 | 1 | 1 |
| occ = 1 (%) | 100 | 100 | 100 | 100 | 100 |
| **Box** | | | | | |
| Lengths (Å) | 248.88, 250.25, 250.25 | 301.12, 301.12, 303.88 | 303.88, 302.50, 301.12 | 333.00, 333.00, 333.00 | 324.50, 324.50, 324.5 |
| Angles (°) | 90.00, 90.00, 90.00 | 90.00, 90.00, 90.00 | 90.00, 90.00, 90.00 | 90.00, 90.00, 90.00 | 90.00, 90.00, 90.00 |
| **Model vs. Data** | | | | | |
| CC (mask) | 0.84 | 0.78 | 0.73 | 0.8 | 0.81 |
| CC (box) | 0.78 | 0.72 | 0.69 | 0.82 | 0.69 |
| CC (peaks) | 0.72 | 0.65 | 0.58 | 0.69 | 0.64 |
| CC (volume) | 0.83 | 0.75 | 0.72 | 0.8 | 0.8 |
| **Resolution range (Å)** | 3.2-4.4 | 3.8-5.8 | 4.1-6.1 | 6.4-8.5 | 3.0-3.6 |

**Table S2. BoxB-like sequence motifs within the NC-3 and NC-4 genomes.**

Genome positions of nucleotide strings fulfilling the search motif together with the color-coded reactivities (black, green, orange and red from low to high) in NC-3 and NC-4. The 13 sequences that show sufficiently low reactivity to potentially act as packaging signals are highlighted in grey. Of these, seven motifs occur in a stem loop in over half of the sample folds for at least 50 *m*,*b* combinations tested. Two are the BoxBr tags (BB1, BB2), introduced by design, and the other five potential packaging signals are designated PS1-PS5 in the order they appear in the sequence.

| URxRxRR | | | |
|---|---|---|---|
| **Position** | **NC-3** | **NC-4** | |
| **32** | UAGAGGG | UAGAGGG | **PS1** |
| **60** | UGAAGAA | UGAAGAA | **BB1** |
| 116 | UAAGAAG | UAAGAAG | |
| **133** | UAUGGGA | UAUGGGA | |
| **135** | UGGGAAA | UGGGAAA | **PS3** |
| **172** | UGAGAAA | UGAGAAA | **PS4** |
| 188 | UGGAAAG | UGGAAAG | |
| 294 | UACGCAA | UACGUAA | |
| **383** | UGGGAAG | UGGGAAG | **PS5** |
| 420 | UAUACAA | UAUACAA | |
| **482** | UACGAAG | UACGAAG | |
| 564 | UGGAGGG | UGGAGGG | |
| 581 | UGCAUAG | UGCAUAG | |
| **604** | UGAAGAA | UGGAGAA | |
| 641 | UGGGAAA | UGGGAAA | |
| 676 | UAAAGAG | UAAAGAG | |
| 734 | UAAGCGG | UAAGCGG | |
| **768** | UGAAGAA | UGAAGAA | **BB2** |

| URxRxxx | | | |
|---|---|---|---|
| **Position** | **NC-3** | **NC-4** | |
| 79 | UAGAGUG | UAGAGUG | |
| **84** | UGUGGGC | UGUGGGC | **PS2** |
| **86** | UGGGCUC | UGGGCUC | |
| 127 | UAUACAU | UAUACAU | |
| 129 | UACAUAU | UACAUAU | |
| 205 | UGGAGCU | UGGAGCU | |
| 211 | UGGAGCA | UGGAGCA | |
| **220** | UGCAAUG | UGCAAUG | |
| 245 | UAUAACG | UAUAACG | |
| 256 | UGUAGUU | UGUAGUU | |
| 288 | UAGAACU | UAGAACU | |
| 298 | CAAACCU | UAAACCU | |
| 322 | UACAGCU | UACAGCU | |
| 336 | UGGAACA | UGGAACA | |
| 406 | UGAAAUG | UGAAAUG | |
| 422 | UACAAGU | UACAAGU | |
| **474** | UUGAAAUC | UUGAAAUC | |
| 490 | UAAACUA | UAAACUA | |
| 631 | UCCAGGC | UCCAGGC | |
| 658 | UGCGGAU | UGCGGAU | |
| 664 | UGAACUG | UGAACUG | |

**Table S3. Pairwise Pearson correlation coefficients (PCCs) for normalized replicates at 50 ms exposure.**

PCCs for triplicate primer extensions, analyzed by primer region and RNA. A, B, and C represent the individual replicates. Lower values imply greater variability in the respective mRNA segment.

| | NC-3 | | | | | NC-4 | | | | |
|---|---|---|---|---|---|---|---|---|---|---|
| | **Transcript** | | | ***In situ*** | | **Transcript** | | | ***In situ*** | |
| **Primer** | **A** | **B** | | **A** | **B** | **A** | **B** | | **A** | **B** |
| **12** | 0.964 | | **B** | 0.718 | | 0.927 | | **B** | 0.899 | |
| | 0.940 | 0.930 | **C** | 0.774 | 0.869 | 0.963 | 0.950 | **C** | 0.893 | 0.976 |
| | **A** | **B** | | **A** | **B** | **A** | **B** | | **A** | **B** |
| **11** | 0.932 | | **B** | 0.919 | | 0.931 | | **B** | 0.975 | |
| | 0.919 | 0.981 | **C** | 0.983 | 0.917 | 0.715 | 0.725 | **C** | 0.949 | 0.954 |
| | **A** | **B** | | **A** | **B** | **A** | **B** | | **A** | **B** |
| **10** | 0.866 | | **B** | 0.865 | | 0.954 | | **B** | 0.952 | |
| | 0.893 | 0.951 | **C** | 0.927 | 0.918 | 0.841 | 0.768 | **C** | 0.957 | 0.954 |
